# The archaeal KEOPS complex possesses a functional Gon7 homolog and has an essential function independent of cellular t^6^A modification level

**DOI:** 10.1101/2022.08.26.505501

**Authors:** Pengju Wu, Qi Gan, Xuemei Zhang, Yunfeng Yang, Yuanxi Xiao, Qunxin She, Jinfeng Ni, Qihong Huang, Yulong Shen

## Abstract

KEOPS is a multi-subunit protein complex conserved in eukaryotes and archaea. It is composed of Pcc1, Kae1, Bud32, Cgi121, and Gon7 in eukaryotes and is primarily involved in N^6^-threonylcarbamoyl adenosine (t^6^A) modification of tRNAs. Recently, KEOPS is reported to participate in homologous recombination repair in yeast. To characterize the KEOPS in archaea (aKEOPS), we conducted genetic and biochemical analyses of its encoding genes in the hyperthermophilic archaeon *Saccharolobus islandicus*. We show that aKEOPS also possesses five subunits, Pcc1, Kae1, Bud32, Cgi121, and Pcc1-like (or Gon7-like), just as eukaryotic KEOPS. Pcc1-like has physical interactions with Kae1 and Pcc1 and can mediate the monomerization of the dimeric subcomplex (Kae1-Pcc1-Pcc1-Kae1), suggesting that Pcc1-like is a functional homolog of the eukaryotic Gon7 subunit. Strikingly, none of the genes encoding aKEOPS subunits, including Pcc1 and Pcc1-like, can be deleted in the wild type and in a t^6^A modification complementary strain named TsaKI, implying that aKEOPS complex is essential for an additional cellular process in this archaeon. Knock-down of the Cgi121 subunit leads to severe growth retardance in the wild type which is partially rescued in TsaKI. These results suggest that aKEOPS plays an essential role independent of cellular t^6^A modification level. In addition, archaeal Cgi121 possesses dsDNA-binding activity which relies on its tRNA 3’ CCA tail binding module. Our study clarifies the subunit organization of archaeal KEOPS and suggests of an origin of eukaryotic Gon7. The study also reveals a possible link between the function in t^6^A modification and the additional function presumably homologous recombination.

## Introduction

KEOPS (Kinase, putative Endopeptidase, and Other Proteins of Small size) is a multi-subunits protein complex conserved in eukaryotes and archaea (1,2). Its subunits Pcc1, Kae1, Bud32, Cgi121 and eukaryotic fifth subunit Gon7 form a linear architecture as Gon7-Pcc1-Kae1-Bud32-Cgi121 or Cgi121-Bud32-Kae1-Pcc1-Pcc1-Kae1-Bud32-Cgi121, in which Pcc1 serves as a dimerization module while Gon7 interacts with Pcc1 to eliminate the Pcc1 dimer (3-5). The primary role of KEOPS is threonylcarbamoyl (TC) modification at A37 of NNU anti-codon tRNAs yielding N^6^-threonylcarbamoyl adenosine (t^6^A) (6). t^6^A modification is critical for stabilizing tRNA anti-codon loop structure and base pairing of the anticodon with the corresponding codon for the enhancement of translation fidelity in all three domains of life (6,7). In humans, defects in t^6^A modification due to Gon7 mutants lead to Galloway-Mowat syndrome, which is characterized by infantile onset of microcephaly and central nervous system abnormalities (8,9).

In eukaryotic cytoplasm and archaea, the KEOPS complex catalyses t^6^A modification in association with Sua5 in two steps. Firstly, Sua5 synthesizes the intermediate TC-AMP and then KEOPS transfers TC to tRNA (10). In the second step, the 3’ CCA ssRNA binding subunit Cgi121 recruits tRNAs to KEOPS complex while Pcc1, Kae1 and Bud32 recognize the NNU tRNA together (11). During TC transfer by KEOPS, the TC transferase Kae1 functions as the active center and the kinase/ATPase subunit Bud32 stimulates the activity of Kae1 (12). However, the precise role of Gon7 in tRNA modification is unclear now.

Besides t^6^A modification, KEOPS also functions in homologous recombination (HR) repair and telomere maintenance in budding yeast, both of which are critical for genome integrity and cell survival (1,13,14). End resection of DNA double-strand breaks (DSB) is an essential step in HR repair (15,16). The Mre11-Rad50-Xrs2/Mre11-Rad50-NBS1(MRX/N) complex in conjunction with Sae2/CtIP initiates the resection which is followed by two distinct pathways mediated by Exo1/EXO1 and Dna2/DNA2-Sgs1/BLM, respectively, generating a long 3’-tail of single-stranded DNA (ssDNA) (17-20). Recent studies reported that in knockout yeast of the KEOPS subunits, the production of ssDNA near DNA break ends and HR efficiency were reduced, indicating that the KEOPS complex is a novel important player in DSB end resection or its regulation (13). Additionally, the deletion of the KEOPS genes in yeast led to telomere shortening, and depletion of KEOPS subunits in the *cdc13-1* mutant strain reduced the amount of telomeric ssDNA, indicating that the complex is also involved in the maintenance of telomeres by regulating telomere replication and recombination (1). Although the mechanism of KEOPS complex in t^6^A modification is relatively clearly known, the precise role of KEOPS complex in DSB repair and telomere maintenance is largely unexplored. Interestingly, the functions of KEOPS in HR and telomere maintenance are independent of the t^6^A modification (13,21).

Archaea harbour eukaryotic-like DNA transaction system and serve as good models for studying the biology of archaea and the evolution of life. In fact, many eukaryotic-like HR proteins of archaea, especially Mre11 and Rad50 and their complexes have served as excellent models for elucidating the structural and functional mechanisms for the eukaryotic counterparts (22-24). In thermophilic archaea, all key proteins involved in HR are essential (25,26), indicating that HR plays fundamental role in DNA repair for the survival of these archaea. Similarly, archaeal KEOPS (aKEOPS) has been used as a model for the structural and biochemical studies of eukaryotic KEOPS (10-12,27). Nevertheless, so far genetic analysis has been studied only in a halophilic euryarchaeon *Haloferax volcanii* for some of the KEOPS subunits (28). Interestingly, in this euryarchaeon, the fused *kae1-bud32* gene is essential, so was the *cgi121*. In contrast, the *pcc1* (encoding the putative dimerizing unit of KEOPS) was not essential. Deletion of the *pcc1* led to pleiotropic phenotypes, including decreased growth rate and increased cellular DNA content (28), implying a failure in chromosome segregation and/or DNA repair. Whether the DSB repair function of KEOPS is conserved or not in archaea and eukaryotes and its relationship with other functions, tRNA modification and telomere maintenance, are unclear up to now.

In this work, we carried out *in vivo* and biochemical analysis of KEOPS subunit in *Saccharolobus islandicus* (formally *Sulfolobus islandicus*) REY15A, a thermophilic model crenarchaeon close to the ancestral eukaryotes. We show that one Pcc1 homolog, Pcc1-like, is the fifth subunit and functional Gon7 homolog of aKEOPS. We further show that each of the five genes encoding aKEOPS subunits (including the gene annotated as *pcc1-like*) is essential in the wild-type and a strain complemented with a bacterial t^6^A modification system, suggesting that KEOPS may have additional and essential functions in *S. islandicus*. Further biochemical analysis suggests that the Pcc1-Kae1 subcomplex and Cgi121 are the main dsDNA binding players, and three key conserved residues of Cgi121 involved in tRNA 3’ CCA tail binding also contribute to dsDNA binding, revealing a possible link between two functions of the complex, t^6^A modification and HR. Finally, a model for the evolutional scenario of t^6^A modification systems is proposed.

## Materials and methods

### Strains and growth conditions

*S. islandicus* strain REY15A (E233S) (Δ*pyrEF*Δ*lacS*) (hereafter E233S) and its transformants were cultured as described previously (29). D-arabinose (0.2 % w/v) was used for cell growth to induce protein overexpression controlled by different *araS* promoters.

### Construction of plasmids and strains

The strains and vectors used and constructed in this study are listed in Supplementary Table S1 and S2. Oligonucleotides for spacers, *P*_*araS*_ mutant promoters, fluorescence-labelled ssDNA, and primers for PCR and RT-qPCR are listed in Supplementary Table S3. For the construction of *Thermotoga maritima* t^6^A biosynthesis system knock-in strain TsaKI, constitutive expression vector pSeGP was constructed by replacing the *araS*-SD sequence with the promoter of *S. islandicus* glutamate dehydrogenase (*P*_*gdhA*_) at the site between SphI and NdeI of pSeSD (30). The gene for the TsaB, TsaC, TsaD, or TsaE was inserted into the NdeI and SalI site of pSeGP individually. Then, the cassettes for the TsaB, TsaC, TsaD, and TsaE were amplified and ligated into pRSF-Duet1 sequentially in the order of TsaB-TsaD-TsaE-TsaC. The homologous arms (L-arm and R-arm) of *amyα* (SiRe_1098) were ligated via gene splicing by overlap extension (SOE) PCR with KpnI/XhoI sites in between and inserted into the SphI/BlnI site of pUC19. Next, the cassette of *Tm*TsaB-TsaD-TsaE-TsaC was inserted into the KpnI/XhoI site between the two homologous arms. Finally, the fragment containing the homologous arms and the *Tm*TsaB-TsaD-TsaE-TsaC was amplified and inserted into the SphI/BlnI site of the pGE vector (31), yielding a vector pGE-*Tm*TsaBDEC*-*KI. To construct pGE-*Tm*TsaD*-*KI, the spacer was designed to target the endo-β-mannanase (SiRe_2238) and the fragment containing the homologous arms and *Tm*TsaD was inserted into the SphI/BlnI site of pGE. For construction of the vectors for the gene knock-down, the corresponding spacers were inserted into the mini-CRISPR locus in pGE (31). For the construction of the knock-out strain, the corresponding spacers were inserted into the mini-CRISPR locus and the fragment containing the homologous arms was inserted into the SphI/XhoI site of pGE. To construct vectors for protein overexpression in *S. islandicus*, the gene fragments were amplified by PCR, digested with restriction enzymes, and inserted into the NdeI and SalI site of pSeSD (32). To construct vectors for protein co-overexpression in *S. islandicus*, another multiple clone site (XhoI/NheI) with an *araS* promoter was introduced into pSeSD yielding pSeSD-2ParaS for plasmid construction.

To construct the knock-in strain TsaKI harbouring *T. maritima* t^6^A biosynthesis system, the vector pGE-*Tm*TsaBDEC-KI was transformed into E233S. The transformants were screened on uracil free media and verified by PCR using flanking primers (Table S3). The vector pGE-*Tm*TsaD-KI was transformed into *amyα::Tm*TsaB/TsaC/TsaE yielding the strain TsaKI containing the whole *T. maritima* t^6^A biosynthesis system. The vectors for knock-down, knock-out, and overexpression of KEOPS genes were transformed into E233S to obtain the corresponding strains.

### Bioinformatic analysis

BLAST and genome context analysis was harnessed to identify Sua5 and KEOPS homolog sequences in archaea and eukaryotes (33). The evolutionary distances were calculated using the *p*-distance method (34). For phylogenetic analysis of Pcc1 and Gon7, 126 homolog sequences from 82 representative species of each phylum of archaea and eukaryotes were aligned by MAFFT v7 (35,36). The evolutionary history was inferred by using the Maximum Likelihood method based on the Le_Gascuel_2008 model (37). Initial tree (s) for the heuristic search were obtained automatically by applying Neighbor-Join and BioNJ algorithms to a matrix of pairwise distances estimated using a JTT model, and then selecting the topology with superior log likelihood value. A discrete Gamma distribution was used to model evolutionary rate differences among sites (5 categories (+G, parameter = 2.8806)). The tree was drawn to scale, with branch lengths measured in the number of substitutions per site. All positions with less than 95% site coverage were eliminated. Evolutionary analyses were conducted in MEGA7 and prettified by iTOL (https://itol.embl.de/) (38).

### Protein expression and purification

For protein expression, the vectors were transformed into *E. coli* BL21(DE3)-Codon plus-RIL strain. No-tagged or hexa-histidine (6×His)-tagged *Sis*Kae1 (Kae1 from *S. islandicus*) and *Sis*Bud32 were expressed using pET15b. His-tagged *Sis*Pcc1-like, *Tko*Cgi121, and *Tko*Cgi121I75E were expressed using pRSF-Duet1. MBP- and His-tagged *Sis*Cgi121 and *Sis*Pcc1 were expressed using pMALc2X. No-tagged *Sis*Kae1 were co-expressed with His-tagged *Sis*Pcc1 using pRSF-Duet1. His-tagged *Sis*Pcc1-like was co-expressed with Flag-tagged *Sis*Kae1 and *Sis*Pcc1 using pRSF-Duet1, respectively. The expression was induced by the addition of 0.1 mM IPTG to 1.0 L of LB media at OD_600_ 0.4-0.6 and the cells were cultivated at 37°C for 4 h. The proteins were purified using nickel affinity chromatography (Ni-NTA) coupled to anion exchange (HiTrap Q FF) and size exclusion chromatography (Superdex 200 Increase 10/300) in a buffer containing 50 mM Tris-HCl pH 8.0, 5 % glycerol, and 50–500 mM NaCl depending on different proteins.

For protein expression and purification in *S. islandicus*, strains carrying the constructed pSeSD-based vectors were cultivated in sucrose medium. Expression was induced by the addition of 0.2 % D-arabinose to 1.0 L cultures at an OD_600_ of 0.2 and the cells were cultured at 75°C for 24 h. His-tagged *Sis*Pcc1 and *Sis*Kae1 were expressed separately. His-tagged *Sis*Bud32 and no tagged *Sis*Kae1 were co-expressed by cloning *kae1-bud32* operon into pSeSD. The subsequent protein purification was the same as described above.

### Pull down assay

His-tag pull down was performed using 10 μM His-*Sis*Pcc1-like protein mixed with 12 μM no-tagged *Sis*Kae1, or 10 μM His-*Sis*Kae1 mixed with 12 μM no-tagged *Sis*Bud32, in 200 μl binding buffer (50 mM NaCl, 50 mM Tris-HCl pH 8.0). The mixture was incubated with 50 μl pre-equilibrated Ni-NTA nickel-charged affinity resin. Binding was allowed to proceed for 10 min at 4°C with mild shaking, followed by three times washes with the wash buffer (binding buffer containing 10 mM imidazole). The resin was resuspended in the elute buffer (binding buffer containing 400 mM imidazole) and the supernatant was analysed by 15 % SDS-PAGE and Coomassie blue staining.

MBP-tag pull down was conducted using 10 μM MBP-*Sis*Cgi121-His mixed with 12 μM His*-Sis*Bud32 in 200 μl binding buffer. The mixture was incubated with 50 μl pre-equilibrated amylose resin. Binding was allowed to proceed for 10 min at 4°C with mild shaking, followed by wash in binding buffer for three times. The resin was resuspended in elute buffer (binding buffer containing 10 mM maltose) and the supernatant was analysed by 15 % SDS-PAGE and Western blotting.

### Size exclusion chromatography (SEC) analysis

SEC analysis of single proteins or combinations of the subunits was conducted using AKTA basic 10 FPLC system (GE healthcare) and a Superdex 200 Increase 10/300 SEC column. Chromatography was performed in a running buffer (200 mM NaCl, 50 mM Tris-HCl pH 8.0, and 5 % glycerol). For complex reconstitution, 30 μM *Sis*Pcc1-like was mixed with 20 μM *Sis*Kae1/*Sis*Pcc1-like, or 6 μM *Sis*Pcc1-like mixed with 30 μM *Sis*Kae1, in 600 μl running buffer at 60°C for 30 min before loading onto the column.

### Growth curve and dot assay

The growth curves of the *S. islandicus* strains were measured as follows. Each strain was pre-cultured in a flask containing 25 ml medium at 75°C for 24 h until the culture reached an OD_600_ of 0.4–0.6. Cells were collected and resuspended in 25 ml ddH_2_O and then inoculated into 30 ml medium to an estimated OD_600_ of 0.04 and cultured at 75°C in a 100 ml flask with shaking. The cell densities were measured at the indicated time intervals. Each strain was analysed in duplicate, and the measurements were performed three times independently. For the growth observation on plates, an overnight-grown culture was diluted to an estimated OD_600_ of 0.2. Cells were diluted in ten-fold gradience and spotted on plates. Plates were incubated at 75°C for 5 days before photos were taken.

### Analysis of t^6^A modification and transcription by RT-qPCR

Bulk tRNA were isolated according to the reported method (39) using hot acid-saturated phenol with slight modification. In particular, the cells were grown in 25 ml medium at 75°C to OD_600_ of 0.8 before collection. The isolated bulk tRNA was analysed according to the reported protocol (40). Briefly, tRNA (5 μg) was boiled for 2 min and cooled down on ice immediately. Then the tRNA were digested with 10 U Nuclease P1 (New England Biolabs, MA, USA) at 37°C overnight followed by dephosphorylation using 10 U bacteria alkaline phosphatase (Thermofisher Scientific, USA) at 65°C for 2 h. The ribonucleotides were analysed by Ultimate 3000 HPLC (Thermofisher Scientific) using Zorbax SB-C18 (250 × 4.6 mm, 5 μM) reverse-phase column (Agilent). Ribonucleotides were separated by linear gradient elution from 2 % to 12.5 % buffer B at a flow rate of 1.5 ml/min. The compositions of the HPLC buffers were as follows: (A) 250 mM NH_4_Ac, pH 6.5; (B) 40 % acetonitrile. The peaks were analysed using the software Chromeleon 7 and the relative t^6^A modification level was described as the ratio t^6^A/Adenosine.

For RT-qPCR analysis, when the cultures were grown to an OD_600_ of 0.8, cells of 2 ml cultures were collected by centrifugation. Total RNA was isolated with 1 ml Trizol (TransGene Biotech, Beijing, China) followed by chloroform and isopropanol extraction. The RNA was precipitated by 75 % ethanol and dissolved in 50 μl DEPC water. RNA (500 ng) was used for cDNA synthesis using an Evo M-ML kit (Accurate Biology, Hunan, China) according to the manufacturer’s instructions. RT-qPCR was conducted by CFX96 Real Time PCR detection system (Bio-Rad) using a SYBR Green Pro Taq HS kit (Accurate Biology) according to the manufacturer’s instructions.

### Electrophoretic mobility shift assay (EMSA)

The EMSA assay was performed in a 20 μl reaction mixture containing 10 nM fluorescence-labelled probes (Supplementary Table S3) and the indicated proteins (0–20 μM) in a binding buffer (50 mM Tris-HCl pH 8.0, 100 μM ATP). The reactions were incubated at 37°C for 30 min. The products were separated by 8 % PAGE in 0.5 × TBE at 4°C. The gels were imaged with Amersham ImageQuant 800 (Cytiva, USA).

## Results

### Pcc1-like is the fifth subunit of KEOPS complex in archaea

Four subunits of KEOPS, Pcc1, Kae1, Bud32, and Cgi121, are highly conserved in archaea and eukaryotes (3). Although the fifth subunit Gon7 were identified in yeast and human (1,2,5), it is still unclear whether aKEOPS contains a counterpart of Gon7. To find a putative Gon7 ortholog in archaea, we searched for the distribution of the homologs of Sua5 and KEOPS subunits in all archaeal and representative eukaryotic phyla. It is confirmed that only Sua5, Pcc1, Kae1, and Bud32, the minimal components for t^6^A modification, are conserved in all archaea and eukaryotes at amino acid sequence level (Fig. 1A). Consistently with the previous report, no sequence homolog of Gon7 is found in archaea as a single domain protein (5). However, in rare cases, in some halophilic archaea Gon7 is present as a domain of a proteasome-activating nucleotidase, such as WP_015321675.1 from *Natronococcus occultus* SP4, WP_007139789.1 from *Halobiforma lacisalsi* AJ5, and ELY65446.1 from *Natronococcus jeotgali* DSM 18795, or MutS (e.g., WP_008364526.1 from *Halorubrum litoreum*) (Fig. 1B). Interestingly, many archaea harbour two Pcc1 paralogs denoted Pcc1 and Pcc1-like, which are encoded by SiRe_1278 and SiRe_1701, respectively, in *Saccharolobus islandicus* REY15A (41). Pcc1 is more widely distributed than Pcc1-like. The distribution of Pcc1-like is also found in most superphyla with the least appearance in the DPANN superphylum (Fig. 1A). Sequence alignment and phylogenetic analysis of Pcc1, Pcc1-like, and Gon7 reveals that Pcc1 exhibits a higher evolution rate than the other subunits (not shown) and archaeal Pcc1-like and eukaryotic Gon7 homologs form a separated lineage which is distal to both archaeal Pcc1 and eukaryotic Pcc1 (Fig. 1B). These results support that one of the two Pcc1 paralogs in archaea, Pcc1-like, is a possible Gon7 homolog of eukaryotes.

**Figure 1.**
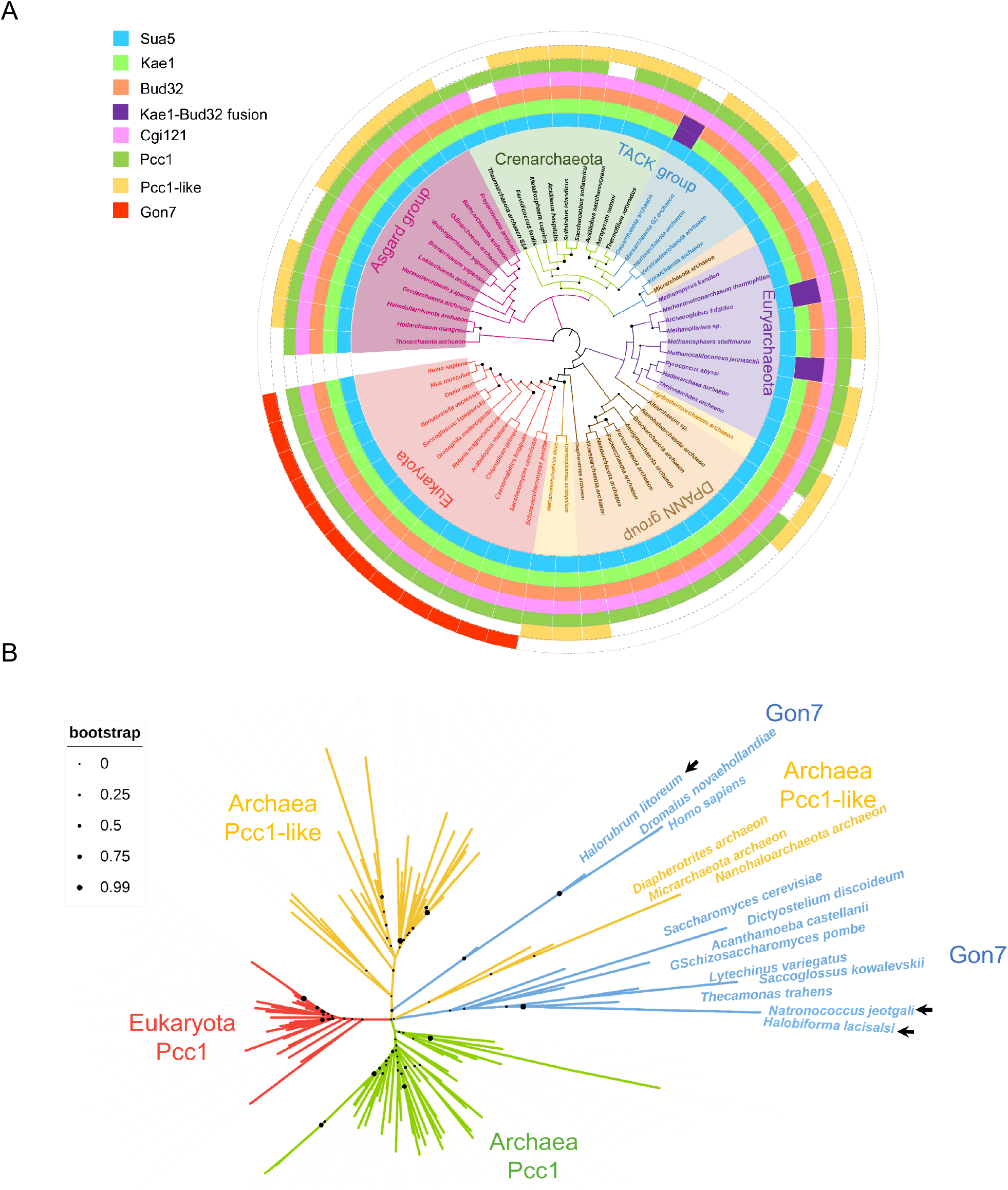
Distribution and phylogenetic analysis of Sua5 and KEOPS subunits in archaea and eukaryotes. (**A**) Distribution of Sua5 and KEOPS subunits. The phylogenetic tree was built based on the amino acid sequences of 59 Kae1 proteins from representative species of each archaea phylum and eukaryotes using the Neighbor-Joining method. The homologs were identified by protein-protein BLAST and protein-nucleotide BLAST in NCBI (https://blast.ncbi.nlm.nih.gov). If no homolog of KEOPS subunits was found by BLAST, genome context analysis was conducted using ribosome genes, which co-occurred with most KEOPS subunits, or XTP dephosphorylase, which co-occurred with the Asgard cgi121, as the references. (**B**) Phylogenetic analysis of Pcc1 and Gon7 homologs. The unrooted maximum likelihood tree of Pcc1, Pcc1-like, and Gon7 homologs was constructed based on 126 sequences from 82 species of archaea and eukaryotes. Filled circles at the nodes indicate bootstrap values of 1000 replicates in different sizes. The arrows indicate Gon7 sequences as a domain of the proteasome-activating nucleotidases (WP_015321675.1 from *Natronococcus occultus* SP4, WP_007139789.1 from *Halobiforma lacisalsi* AJ5, and ELY65446.1 from *Natronococcus jeotgali* DSM 18795), or MutS (WP_008364526.1 from *Halorubrum litoreum*) of halobacteria of euryarchaea. The sequences are available upon request.

To further confirm whether Pcc1-like is involved in the formation of aKEOPS *in vivo*, we conducted affinity purification and mass spectrometry analysis by overexpression of the tagged subunits in *S. islandicus* REY15A. As shown in Table 1, both Pcc1 and Pcc1-like were co-purified with Kae1 and Kae1-Bud32 from the Kae1 and Kae1-Bud32 overexpression cells, respectively, while Kae1 and Pcc1-like were co-purified with Pcc1 from the Pcc1 overexpression cells. Intriguingly, Sua5, which catalyses the synthesis of t^6^A intermediate TC-AMP, was also co-purified with Kae1 and Kae1-Bud32 from their overexpression cells, respectively, indicating that Sua5 might interact with KEOPS complex *in vivo*. Next, *in vitro* pull down analysis showed that Pcc1-like was able to pull down Pcc1 and Kae1, respectively (Fig. S1A). Therefore, we provide strong evidence suggesting that Pcc1-like was the fifth subunit of KEOPS in Sulfolobales.

**Table 1.**
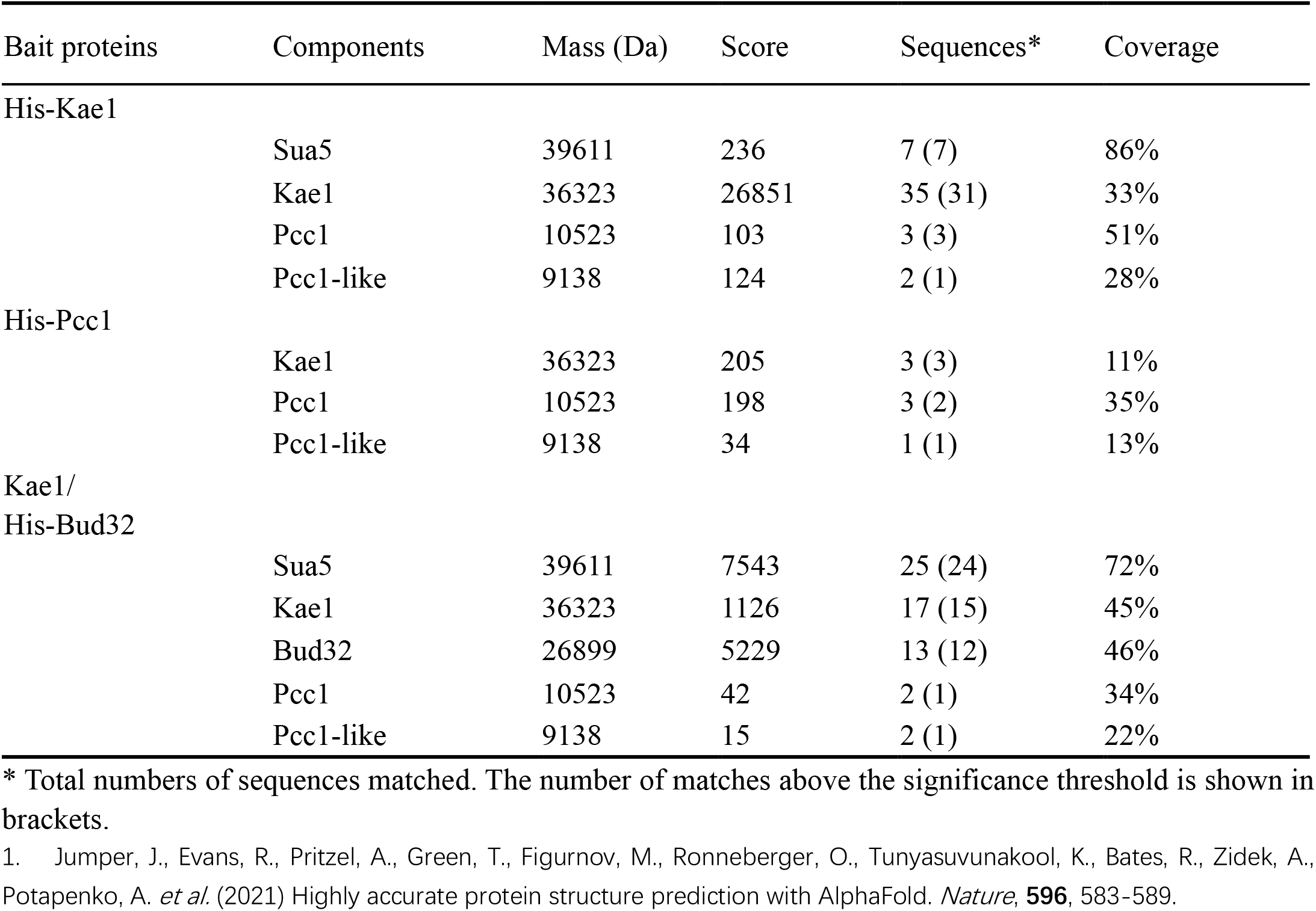
Sua5 and KEOPS subunits identified from co-purified proteins of various KEOPS subunits expressed in *S. islandicus* by mass spectrometry.

### Pcc1-like eliminates Pcc1-mediated dimerization of Pcc1-Kae1 subcomplex *in vitro*

A linear four subunits organization of Pcc1:Kae1:Bud32:Cgi121 was proposed for the KEOPS complex of a euryarchaeon *Methanococcus jannaschii* and budding yeast *S. cerevisiae* (3,4). Thus far, there is no study on the subunit interaction of KEOPS from crenarchaea. To determine the inter-subunit relationship of the crenarchaeal KEOPS, each subunit (except for Pcc1) of the KEOPS from *S. islandicus* was expressed in and purified from *E. coli* and pull-down assay was performed. As shown in Fig. S1B and S1C, Bud32 was pulled down by His-tagged Kae1 and MBP-tagged Cgi121, respectively, verifying that there are interactions between Bud32 and Kae1 as well as between Bud32 and Cgi121. Since Pcc1 is insoluble when it is expressed in *E. coli* alone, we co-express Pcc1 and Kae1 in *E. coli*. His-Pcc1 is able to pull down Kae1, confirming that Pcc1 interacts with Kae1 (Fig. S1A, middle). Therefore, linear subunit organization of Pcc1:Kae1:Bud32:Cgi121 should also be the case for KEOPS from crenarchaea.

However, our analysis of the interaction between Kae1 and Pcc1-like indicate that the organization of Pcc1, Pcc1-like, and Kae1 from *S. islandicus* was not in a linear mode. It was reported that *M. jannaschii* and *S. cerevisiae* KEOPS complex exists as a dimeric structure mediated by a Pcc1 dimer, each monomer of which interacts with a separate Kae1 molecule (27). The fifth subunit of Eukaryotic KEOPS Gon7 interacts with Pcc1 and abolishes this dimerization (4). To understand whether Pcc1-like functions as Pcc1 or an evolutionary intermediate corresponding to Gon7, we examined subcomplex formation among Pcc1, Pcc1-like, and Kae1 by size exclusion chromatography (SEC). As shown in Fig. 2A, the peak of co-expressed Kae1-Pcc1 subcomplex is corresponding to a 90 kDa complex, which was close to a 2:2 subcomplex with the expected molecular mass of 96 kDa. Co-expressed Kae1-Pcc1-like subcomplex was eluted at a peak of 66 kDa corresponding to a Kae1:2 Pcc1-like subcomplex with the predicted molecular mass of 59 kDa when Pcc1-like was about 10 folds over Kae1 (Fig. 2B), indicating that Pcc1-like could not mediate the dimerization of Kae1-Pcc1-like subcomplex (ca. 96 kDa). When Kae1 was excessed (5 folds of Pcc1-like), the stoichiometry of Kae1-Pcc1-like subcomplex became 1:1 with an observed mass of 42 kDa close to the expected mass of 48 kDa (Fig. 2B). Notably, when 1.5 folds Pcc1-like was added into the mixture of 2Kae1:2Pcc1 subcomplex, the 2:2 peak disappeared and a 51 kDa peak was formed (Fig. 2A), suggesting a 1:1:1 stoichiometry of Kae1:Pcc1:Pcc1-like. It should be noted that the elution volume of Kae1:Pcc1:Pcc1-like subcomplex was between those of Kae1:2 Pcc1-like and Kae1:Pcc1-like, but not close to that of Kae1:2 Pcc1-like, suggesting that there was a putative conformational change after the ternary subcomplex was formed. These results suggest that Pcc1 mediates the dimerization of Kae1-Pcc1 while Pcc1-like disrupts the 2Kae1:2Pcc1 subcomplex and stabilizes the Kae1-Pcc1 subcomplex, the same as eukaryotic Pcc1 and Gon7 in eukaryotes (4,11). In *M. jannaschii*, there was a small conformational change when the 2:2 Kae1-Pcc1 subcomplex changed to a 1:2 one, implying a transition state during Pcc1-like evolution (27). Our structural superposition shows that Pcc1-like matches with yeast and human Gon7 proteins with two β sheets and one α helix at the interaction surface of Gon7-Pcc1 (Fig. 2C)(42), reinforcing that Pcc1-like is a potential Gon7 homolog. Hence, we propose that archaea Pcc1-like is the putative functional and structural homolog of Gon7.

**Figure 2.**
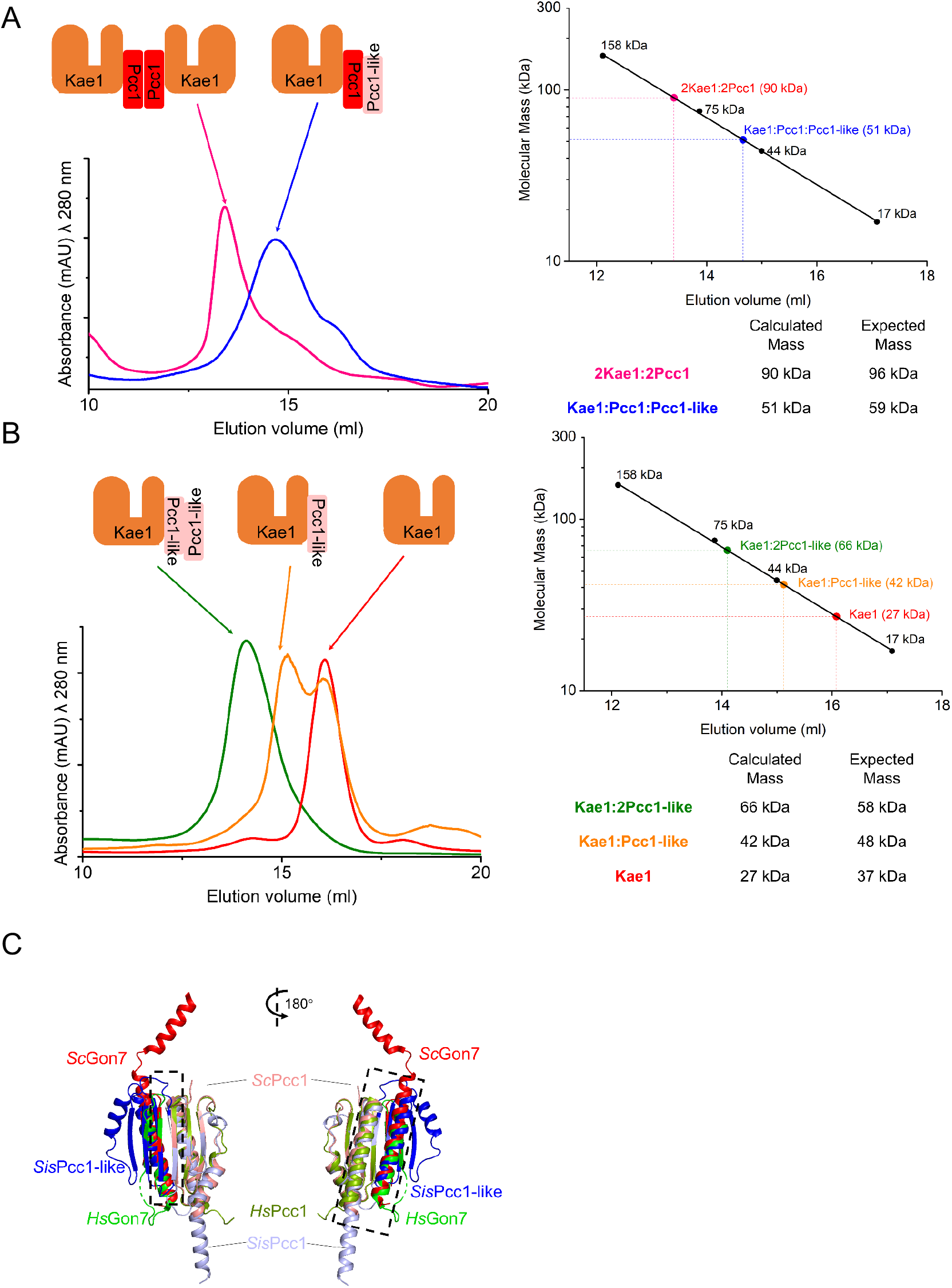
Subunit interaction and complex formation of Pcc1, Pcc1-like, and Kae1 revealed by pull down and size exclusion chromatography (SEC). (**A**) SEC profiles of Kae1/Pcc1 (pink) and Kae1/Pcc1+Pcc1-like (blue). (**B**) SEC profiles of Kae1/Pcc1-like (Pcc1-like 10-fold excessed, green), Kae1/Pcc1-like (Kae1 5-fold excessed, orange), and Kae1 (red). The molecular masses of subcomplexes were calculated based on the protein standards (**A** and **B**, right). (**C**) Structural modeling of *S. islandicus* (*Sis*) Pcc1:Pcc1-like and its superposition with *Saccharomyces cerevisiae* (Sc) Pcc1:Gon7 and *Homo sapiens* (*Hs)* Pcc1:Gon7. The modeling was performed using the online tool AlphaFold2_advanced (https://colab.research.google.com/github/sokrypton/ColabFold/blob/main/beta/AlphaFold2_advanced.ipynb) based on the structures of the *Pyrococcus furiosus* homologs. Structure alignment was conducted using the cealign (CE algorithm) in PyMOL.

### All the genes encoding aKEOPS subunits are essential for cell viability in *S. islandicus*

In eukaryotes, KEOPS functions in t^6^A modification, homologous recombination (HR) repair and telomere maintenance, while only the t^6^A modification function of KEOPS was found in archaea (10,13,21). Telomeres were absent in archaea probably due to their circular chromosomes (43). Therefore, whether aKEOPS also functions in HR repair is important to understand the original functions and mechanisms of KEOPS. To address the role of aKEOPS, we attempted to knock out each of its encoding genes in the crenarchaeon using the genome-editing method based on the endogenous CRISPR-Cas systems (31). These included the genes coding for Pcc1-like, Pcc1, Kae1, Bud32, and Cgi121. We found that none of these genes could be deleted after more than three attempts (Fig. S2), consistent with a previous report that they are essential in the *S. islandicus* M16.4 strain (44). In bacteria, the genes involved in t^6^A synthesis could not be knocked out in nearly all species, possibly due to their essential role in the t^6^A modification (45). In budding yeast, deletion of the KEOPS subunit genes resulted in a t^6^A modification defect and severe slow growth (46). Therefore, this raised a question if the observed essentiality of KEOPS genes in *S. islandicus* could be attributed to the t^6^A modification function or other fundamental ones that are essential for cell viability such as HR repair in Sulfolobales (25,26).

### Knock-in of the *Thermotoga maritima* t^6^A modification system into *S. islandicus* genome

In bacteria, TsaC (Sua5 homolog in bacteria) and the threonylcarbamoyl (TC) transferase complex TCT complex (TsaB/TsaD/TsaE) function in the t^6^A biosynthesis (47-49). To complement the t^6^A modification role of aKEOPS in *S. islandicus*, we first tested if the bacterial t^6^A modification system could be expressed in *S. islandicus* by construction of strains overexpressing each of the genes of the t^6^A modification system from the hyperthermophilic bacteria *Thermotoga maritima* (47). Expression of each gene of the bacterial t^6^A modification system was verified by Western blotting using antibody against the hexa-histidine tag (Fig. S3A). Then a knock-in plasmid was constructed carrying *Tm*TsaB/TsaD/TsaE/TsaC, a tandem array of four genes, each of which was fused to the constitutive promoter of glutamate dehydrogenase (*P*_*gdhA*_) of *S. islandicus* and expressed without the His-tag. After transformation and colony screening, genotypes having only two and three of the genes, *amyα::Tm*TsaE/TsaC and *Tm*TsaB/TsaE/TsaC, respectively, were obtained, suggesting recombination between their *P*_*gdhA*_ promoters (586 bp) could have occurred during the transformation and gene editing processes (Fig. S3B and S3C). Then, the fourth gene *Tm*TsaD was inserted into a gene encoding the endo-β-mannanase in the *amyα::Tm*TsaB/TsaE/TsaC strain, yielding a strain containing the complete *T. maritima* t^6^A modification system (named as TsaKI) (Fig. 3A). However, the t^6^A level in TsaKI was similar to, rather than significantly higher than, that in E233S (Fig. 4B). Considering that each subunit is able to be expressed individually, and the bacterial complex can be formed using individually expressed and purified proteins (47), we assume that the introduced bacterial t^6^A modification system is functional in *S. islandicus*. No increase of t^6^A level in TsaKI is possibly due to that the t^6^A level is already at the upper limit in E233S. The growth of TsaKI was slightly slower than the wild-type E233S (Fig. 3B). It is not clear whether this could be due to disruption of the endo-β-mannanase gene now.

**Figure 3.**
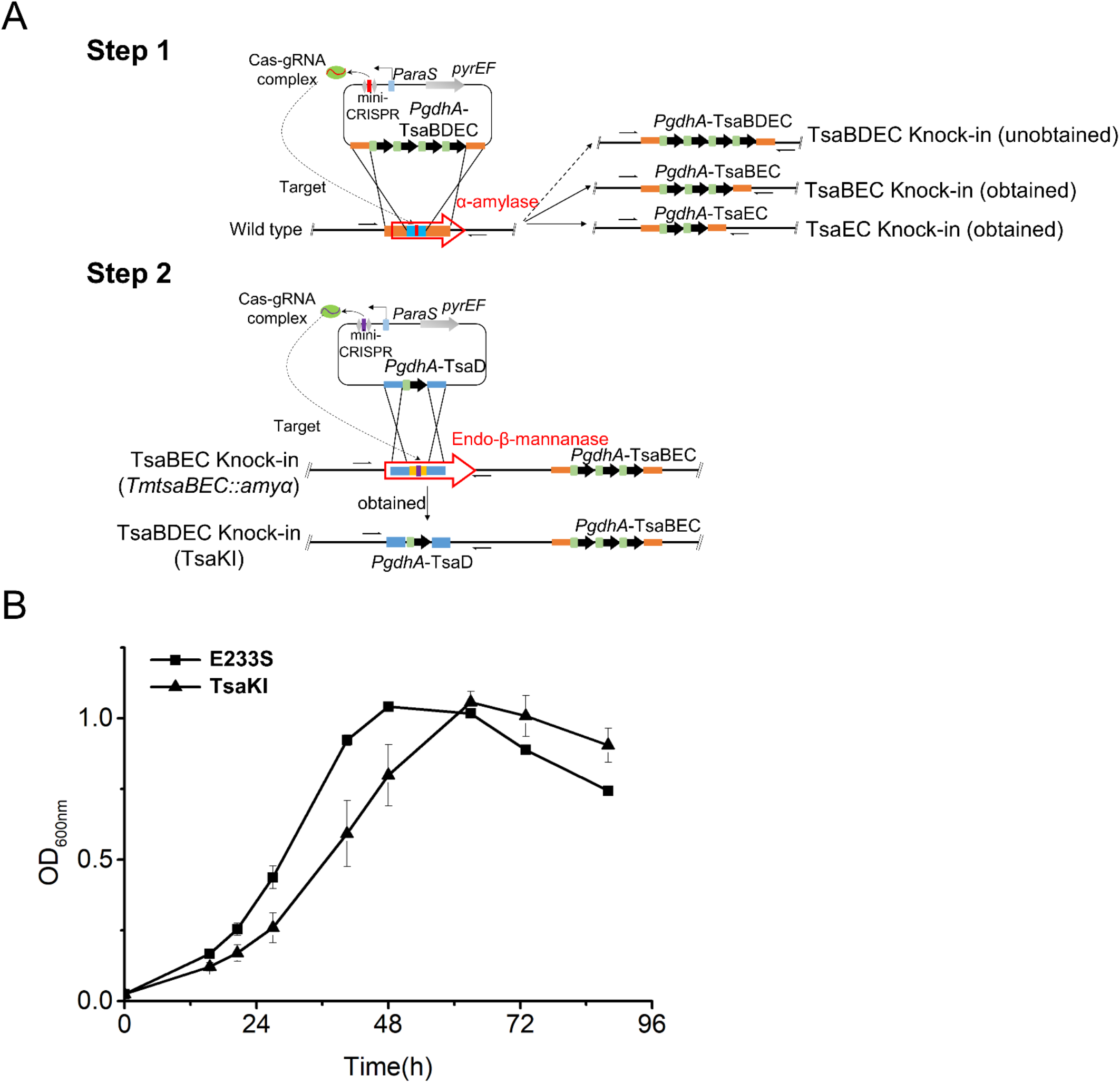
Construction of knock-in strain TsaKI harboring the *T. maritima* t^6^A modification system. (**A**) Schematic for the construction of TsaKI. Step 1, plasmid was constructed to knock in all the four genes of the subunits of *Tm* t^6^A modification system, *TsaB/D/E/C*, at the *α-amylase* locus of *S. islandicus* genome using the endogenous CRISPR-Cas-based method. Strains containing three (*TsaB/E/C*) and two (*TsaE/C*) genes of the subunits were obtained. Step 2, *TsaD* was knocked in at the locus coding for the β-mannanase using the obtained *Tm*TsaBEC knock-in strain, yielding TsaKI. Green rectangle, the promoter of *gdhA* (*PgdhA*). Black arrow, the genes of *TsaB/C/D/E*. (**B**) Growth curves of the wild type (E233S, filled square) and TsaKI (filled triangle). The growth curves were draw based on averages of OD values at 600 nm of three independent cultures. Error bars indicate standard deviations.

**Figure 4.**
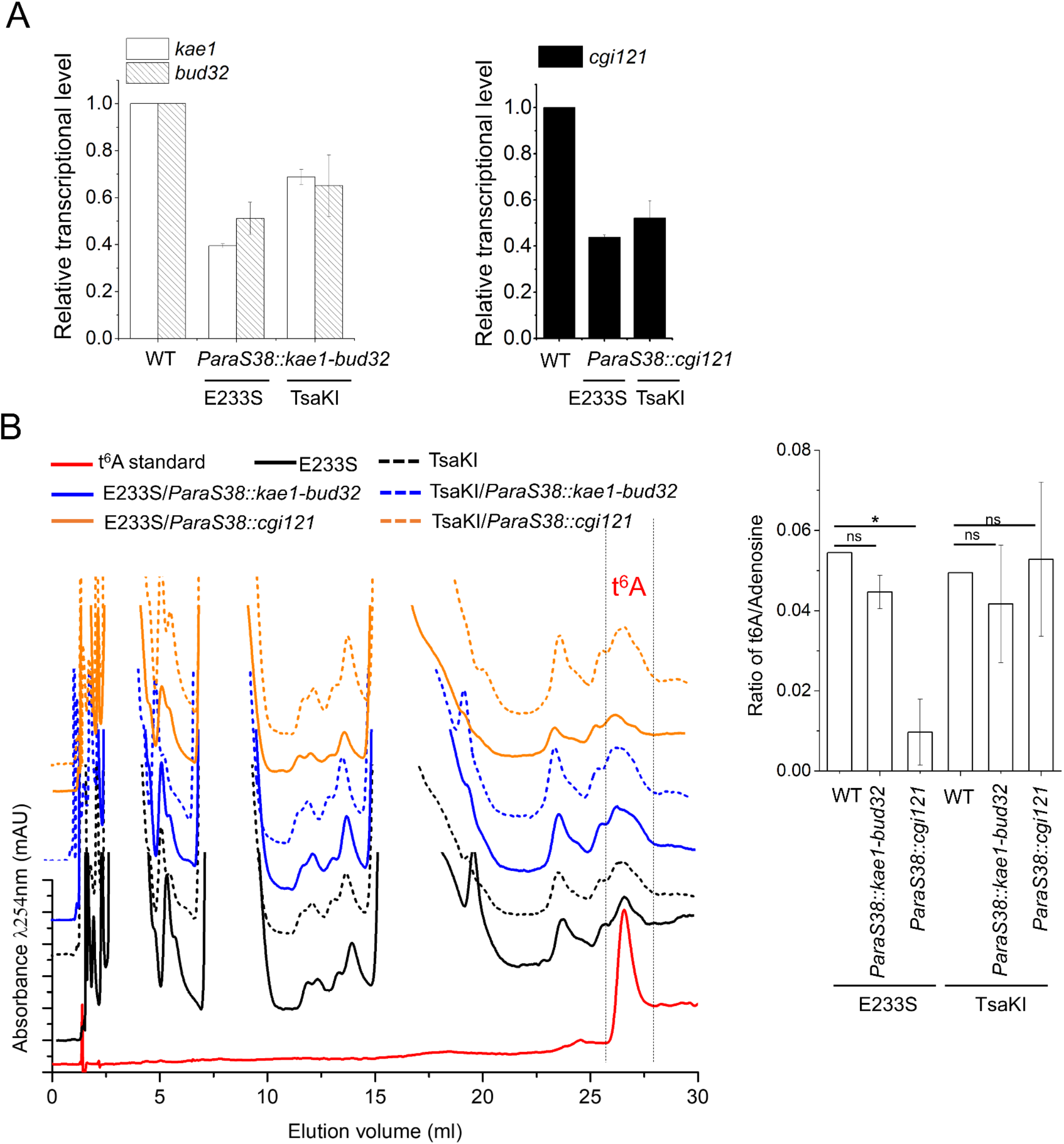
Knock-down analysis of genes encoding *Sis*KEOPS subunits in E233S and TsaKI. (**A**) RT-qPCR of *kae1-bud32* and *cgi121* knock-down strains with (TsaKI) and without the bacterial system (E233S). (**B**) Representative HPLC profiles of nucleoside composition from each sample (left) and quantification (right) showing the average t^6^A content normalized to the content of adenosine (n = 2 independent experiment samples, ±SD). The samples were from the wild type strain E233S (black), TsaKI (dashed black), *kae1-bud32* knock-down in E233S (blue), *cgi121* knock-down in E233S (orange), *kae1-bud32* knock-down in TsaKI (dashed blue), and *cgi121* knock-down in TsaKI (dashed orange), respectively. The t^6^A standard is shown in red. The t^6^A levels were calculated as average ratio of t^6^A/Adenosine. Significance: ns, not significant; *, p <0.05.

### The bacterial t^6^A modification system complements t^6^A modification defect in Cgi121 knock-down mutant

To analyse whether aKEOPS functions beyond t^6^A modification, the KEOPS knock-out plasmids were introduced into TsaKI along with E233S again. We found that similar transformation frequencies were obtained for all KEOPS knock-out plasmids in both strains, which was much lower than that of pSeSD, a reference plasmid, suggesting that these transformants could carry escape mutations that inactivate the CRISPR immune responses. Indeed, analysis of their genotypes revealed that all transformants were wild type (data not shown), indicating that KEOPS subunits have other essential function *in vivo* in addition to the t^6^A modification.

Since all KEOPS subunits are essential either in E233S or TsaKI, we performed gene knock-down analysis to identify the essential function in *S. islandicus*. A previous transcriptomic study showed that the transcripts of *pcc1, pcc1-like, cgi121*, and *bud32* were quite low (FPKM~60-140), while those of *kae1* were at a moderate level (FPKM~800-1000) (50). In addition, CRISPR knock-down in this archaeon is based on Type III-B CRISPR-Cas systems, and a prerequisite for the gene knock-down experiment is the presence of a protospacer-flanking sequence in target genes matching 5’-GAAAG-3’, the 5’-handle tag derived from the CRISPR repeat sequences (41). However, such a motif is lacking in the coding sequence of *cgi121*. Therefore, we harnessed two methods for gene knock-down: CRISPR-Cas-based mRNA cleavage for *pcc1, pcc1-like, kae1*, and *bud32*, and replacement of the native promoters with mutant *P*_*araS*_ (*P*_*araS*_*24* or *P*_*araS*_*38*) (51) for *cgi121* and the *kae1-bud32* operon (Fig. S4A and S4B). Proteins can only be slightly expressed in the presence of sucrose and induced (about 20-fold) in the presence of arabinose with the mutant promoters, and the expression level with *P*_*araS*_*38* mutant is expected to be a quarter of that with *P*_*araS*_*24* in sucrose (51).

As shown in Fig. 4A, the transcriptional levels of *kae1, bud32*, and *cgi121* in the *P*_*araS*_*38* (left and middle) replacement strains were reduced to 40-70 % of those of the wild type in E233S and TsaKI in sucrose medium.

To study whether the t^6^A modification level was affected in the knock-down strains, we estimated their t^6^A modification levels by HPLC. Knock-down of *kae1-bud32* in E233S or TsaKI did not lead to any significant difference in t^6^A levels as compared with the control (Fig. 4B). In contrast, the *cgi121* knock-down led to a dramatic reduction (p-value=0.029) in t^6^A modification level in the knock-down E233S strain. However, the t^6^A modification level in the *cgi121* knock-down TsaKI strain did not change significantly as compared with that in E233S (Fig. 4B). Since the mRNA level of *cgi121* knock-down in E233S was 0.44±0.01 of that in wild type, close to the *cgi121* knock-down ratio in TsaKI (0.52±0.07). Therefore, the relatively higher t^6^A levels in TsaKI should not be due to Cgi121 knock-down efficiency but due to introduction of the Tsa system. The results indicate that Cgi121 is critical for t^6^A modification in E233S. The fact that knock-down of *cgi121* in TsaKI strain did not affect the t^6^A modification level supports that the bacterial t^6^A modification system complements reduction of t^6^A modification level of *S. islandicus* KEOPS.

### The *cgi121* knock-down strains display growth retardance

To reveal the additional function of aKEOPS, we analyzed the growth of the *cgi121* knock-down and the *kae1-bud32* knock-down strains in liquid medium and on plates. As shown in Fig. 5A, while *kae1-bud32* knock-down E233S did not show any difference compared with the wild type E233S, the *cgi121* knock-down E233S strain displayed severe growth retardation in sucrose medium. Interestingly, the *cgi121* knock-down TasKI strain grew better than the *cgi121* knock-down E233S, although it still grew apparently slower than E233S and TsaKI, while the *kae1-bud32* knock-down TsaKI grew slightly faster than TasKI. On plates, the growth differences of the strains were similar to those in the liquid (Fig. 5B). For example, the wild type E233S was able to grow at the dilutions from 10^0^ to 10^−5^, but *cgi121* knock-down E233S only grew at 10^0^ dilution (Fig. 5A). The *cgi121* knock-down TsaKI grew at 10^0^ -10^−2^ dilutions not fully rescued, consistent with the growth curves (Fig. 5A and 5B). Intriguingly, the growth differences disappeared when the strains were cultivated in medium containing arabinose (Fig. 5A). The results indicate that KEOPS has an additional function independent of t^6^A modification level which is related to cell growth.

**Figure 5.**
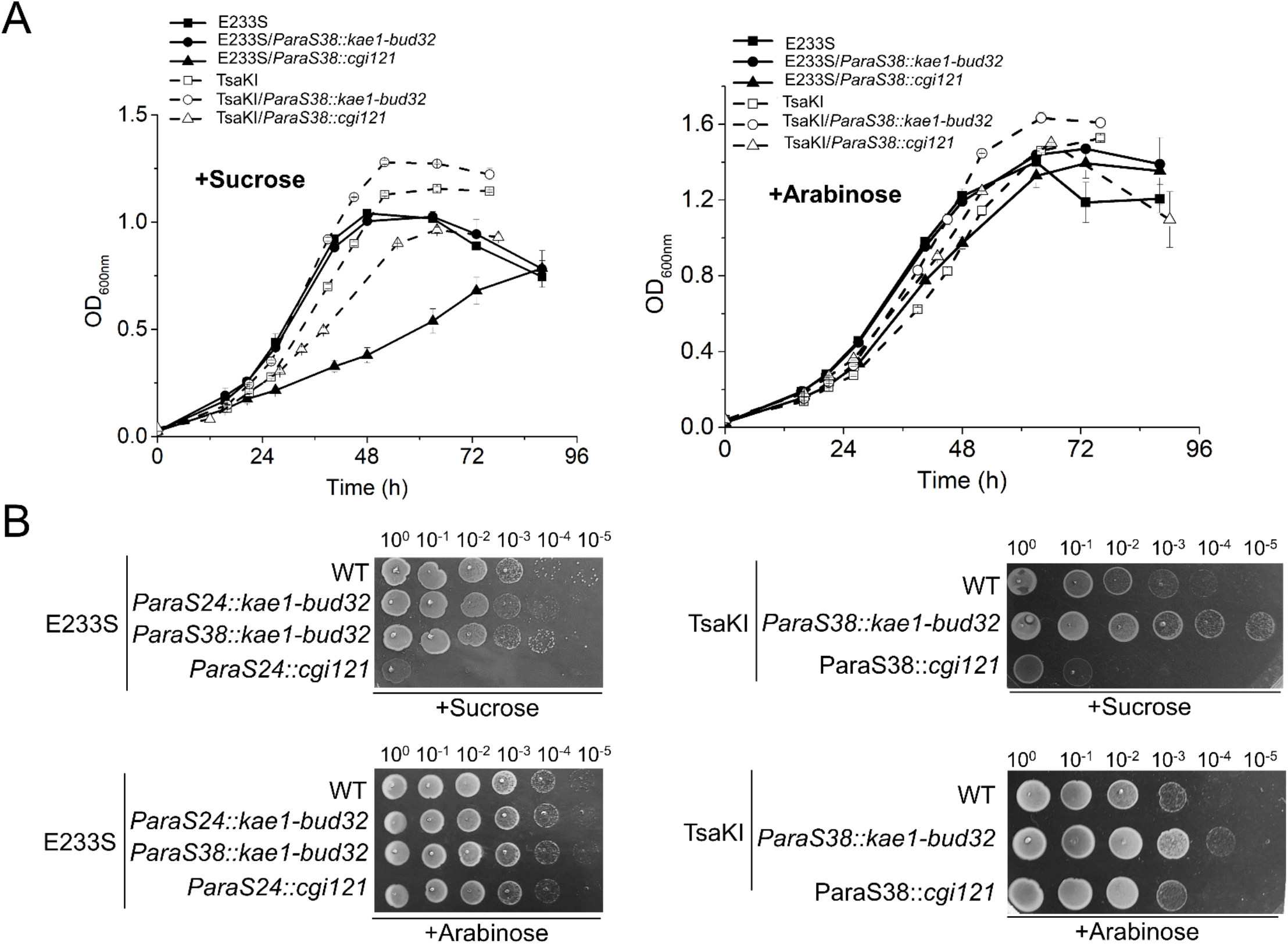
Growth analysis of *kae1-bud32* and *cgi121* knock-down strains in E233S and TsaKI. **(A)** Liquid growth. The OD values at 600 nm were measured every 6 or 12 h. The growth curves were draw based on averages of three independent cultures. Error bars indicate standard deviations. The media containing sucrose (left) and arabinose (right) were used. **(B)** Spot assay using 10 μl of a series of 10-fold dilutions prepared from a starting suspension of an OD_600_ of 0.2. The samples were spotted on the plates with media containing sucrose (above) or arabinose (bottom). The plates were cultured at 75 °C for 5 days before photographing.

Although the transcriptional level of *kae1*-*bud32* decreased in the *kae1-bud32* knock-down strain, there is no apparent difference in growth between the wild type and *kae1-bud32* knock-down strains (Fig. 5). This may be due to relatively high levels of Kae1 and Bud32 in the cells. The amount of functional KEOPS complex might be limited by the subunit with a lowest amount in the cell which might be Cgi121, and the knock-down of *kae1-bud32* could not affect the formation of functional KEOPS complex. The exact mechanism needs further investigation.

### Cgi121 exhibits dsDNA binding activity

In t^6^A modification, tRNA recruitment and recognition is the key step, in which Cgi121 is responsible for tRNA recruitment while other subunits, Pcc1, Kae1, and Bud32, contribute to specific tRNA recognition (11). Interestingly, reconstructed *S. cerevisiae* KEOPS complex exhibits ssDNA and dsDNA binding activity *in vitro* (13). Hence, we propose a hypothesis that there might be a similar nucleic acid binding mechanism of KEOPS functions shared in DNA repair and t^6^A modification. To reveal this, we analysed the DNA binding activity of Cgi121 and other subunits and subcomplexes. We found that Cgi121 and Bud32 were able to bind to long dsDNA (Fig. S5A), although the binding activity of Bud32 was much weaker than that of Cgi121. Intriguingly, Kae1 and Pcc1 in combination, not Kae1 or Pcc1 alone, also exhibited dsDNA binding activity (Fig. S5B).

It was shown that *M. jannaschii* Cgi121 is an ssRNA binding protein which can specifically recognize tRNA 3’ CCA tail (11). Since the protein was always degraded during protein expression and purification using a 6×His tag, a maltose-binding protein (MBP) tag (40 kDa) was fused to *Sis*Cgi121 for the purification. To exclude the potential effect of the MBP tag on DNA binding ability of *Sis*Cgi121, we also purified the euryarchaeal *T. kodakarensis* Cgi121 (*Tko*Cgi121), which has about 21 % sequence identity to *Mj*Cgi121 and 17 % to *Sis*Cgi121. Meanwhile, the tRNA 3’ CCA tail binding deficient mutant K59E, Q74A and I75E of *Tko*Cgi121 were also constructed to explore the relationship between dsDNA binding and tRNA recruitment of Cgi121 (Fig. 6A). MBP-tagged SisCgi121 was able to bind dsDNA in relative low concentration although at high concentration (25 μM) the MBP tag also bound to dsDNA (Fig. 6B). Interestingly, the wild type *Tko*Cgi121 was able to bind to dsDNA while the K59E and I75E mutant lost the binding activity (Fig. 6B). Notably, Q74A mutant exhibited partial lose in its dsDNA affinity. Similar result was reported previously for t^6^A modification by *Mj*Cgi121 mutants (11), suggesting a possible same mechanism between dsDNA and ssRNA binding of Cgi121. To investigate whether Cgi121 only binds CCA motif on DNA, 5’-FAM labeled ssDNA and dsDNA with two CCA motifs (45 nt) were analysed. However, all substrates cannot be bound by the wild type *Tko*Cgi121 except for I75E mutant and SisCgi121 at concentration of 20 μM, which is likely nonspecific binding (Fig. S5C and S5D). These results reveal that the tRNA binding residues of *Tko*Cgi121 also contribute to its long dsDNA binding activity, but not short ss/dsDNA binding, suggesting that the putative DNA repair function of KEOPS is related to its t^6^A modification mechanism.

**Figure 6.**
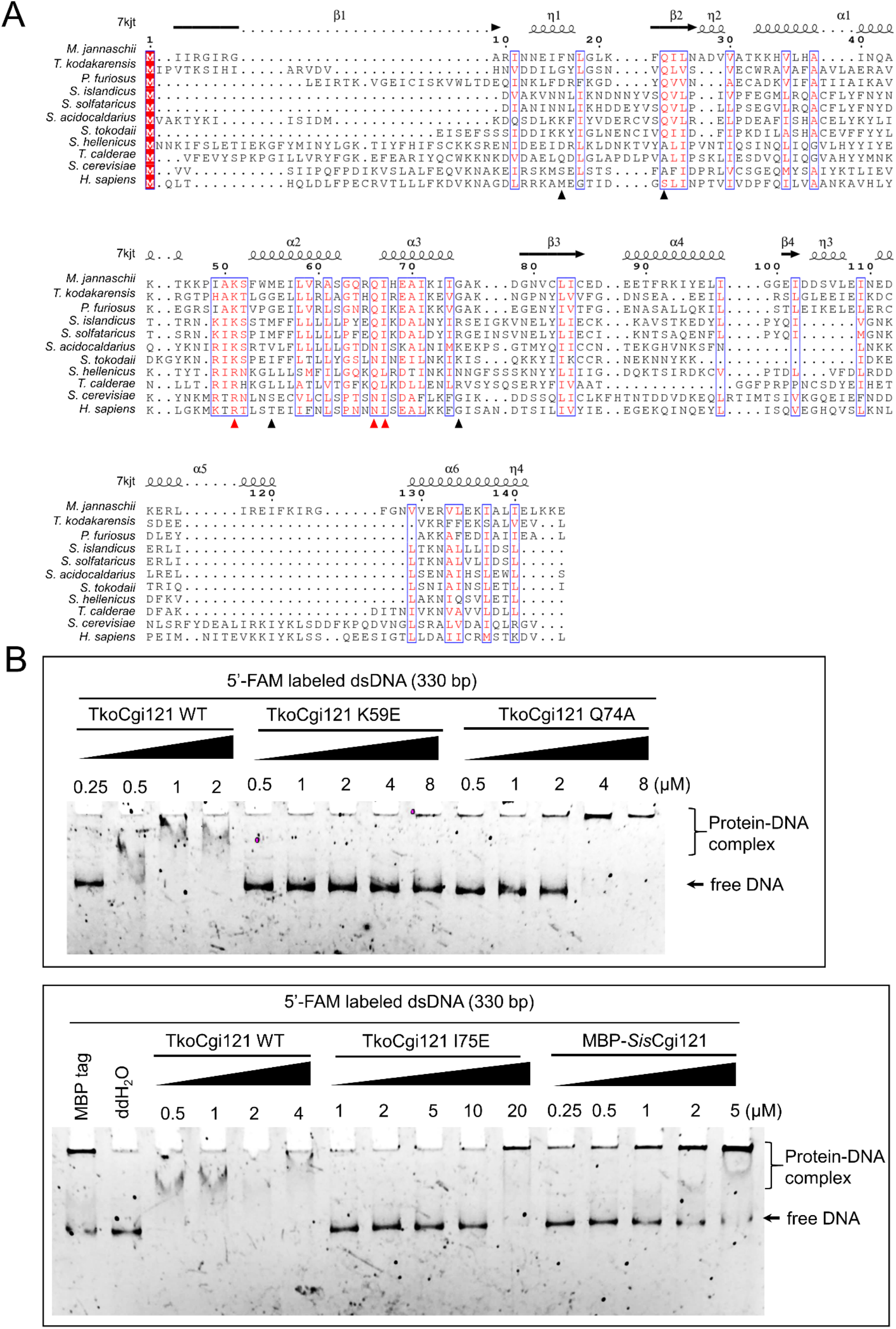
Analysis of the DNA binding ability of wild type Cgi121 and its mutants. (**A**) Structural based sequence alignment of Cgi121. Sequence alignment was executed on MAFFT (v7.505) and modified by Escript 3.0. The residues of the 3’ CCA tail ssRNA binding module of Cgi121 were marked with solid triangles and the most conserved three were colored in red. (**B**) Binding of different Cgi121 proteins for dsDNA (330 bp random sequence). FAM-labeled dsDNA (1 nM) was incubated with samples of the indicated concentrations of Cgi121 at 37°C for 30 min. Native PAGE gels were used for the analysis. The gels were scanned using Amersham ImageQuant 800 after electrophoresis. The detailed reaction conditions are described in the Materials and Methods.

### Overexpression of KEOPS subunits in *S. islandicus*

To further investigate the role of KEOPS subunits in DNA repair and to get insight into the involved mechanisms, the subunits were overexpressed alone or in different combinations in *S. islandicus*. We found that overexpression of either Kae1 or Bud32, the two core components of KEOPS complex, inhibited the cell growth, with the latter showing more profound growth inhibition than the former (Fig. 7A). Interestingly, overexpression of a kinase/ATPase inactive mutant Bud32D134A did not have this inhibition, suggesting that the growth retardation was due to its kinase/ATPase activity (Fig. 7A). When Kae1 and Bud32 were co-overexpressed in a native gene organization manner (Fig. 7C), no growth retardation was observed. However, cell growth was inhibited if the expression of the two genes was controlled by two separate *P*_*araS*_ promoters and induced by arabinose (Fig. 7A and 7C). In a previous study, it was shown that the transcript of *bud32* was several folds lower than that of *kae1* (50). This could be attributed to the difference in the stability of the two transcripts. Alternatively, an unknown mechanism could exist to maintain the ratio of Kae1 and Bud32 *in vivo*. Additionally, we also analysed the effect of the expression of Pcc1, Pcc1-like, and Cgi121, and co-expression of Cgi121 and Bud32 on the cell growth. When overexpressed alone in *S. islandicus*, the proteins of Pcc1, Pcc1-like, and Cgi121 could not be detected by Western blotting. However, when Cgi121 and Bud32 were co-expressed, both proteins were clearly detected (Fig. 7B), indicating that Cgi121 and Bud32 may stabilize with each other *in vivo*. Interestingly, the growth of Cgi121 and Bud32 co-overexpression strain had no difference with the strain carrying the control plasmid pSeSD. Taken together, our results imply that Kae1 and Bud32 and the kinase/ATPase activity of Bud32 have significant physiological functions and that Kae1 and Cgi121 have regulatory roles for the activity of Bud32. The results also suggest that the levels of KEOPS subunits, especially Bud32, Kae1, and Cgi121 must be maintained in an appropriate proportion in *S. islandicus* for their stability and proper functions.

**Figure 7.**
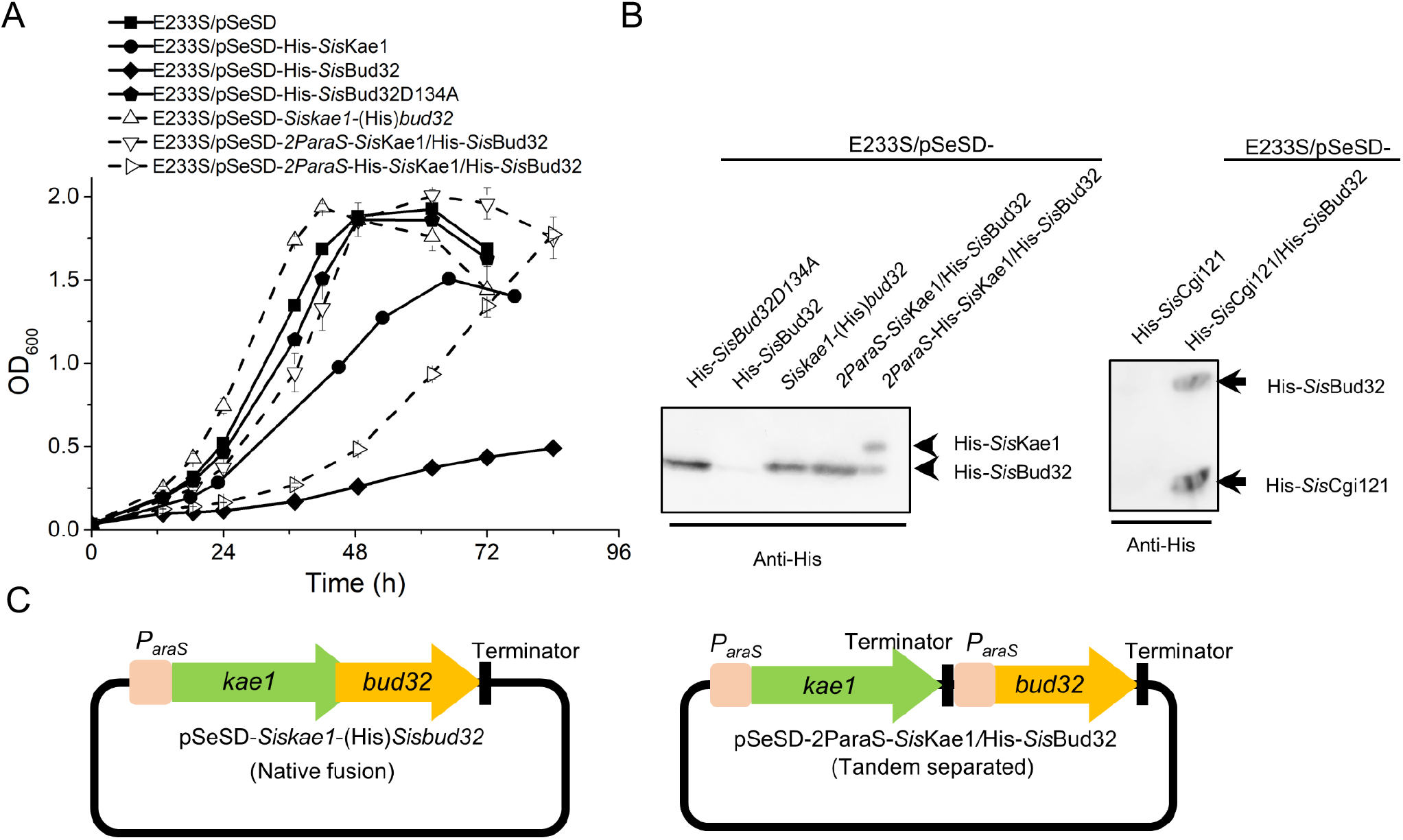
Effects of the overexpression of the KEOPS subunit on cell growth. (**A**) Overexpression of Kae1 and/or Bud32. The overexpression strains were cultured in arabinose medium at an initial OD_600_~0.05. The OD values were measured every 6 or 12 h. The growth curves were draw based on the averages of three independent cultures. Error bars indicate standard deviation. E233S/pSeSD was used as a control. (**B**) Western blotting analysis of the expression of Kae1, Bud32, and Cgi121. The cells were collected at the exponential phase and lysed by boiling for 10 min. The protein levels were detected using anti-His antibody. (**C**) Schematic of the constructs for *Sis*Kae1 and *Sis*Bud32 co-expression. *Kae1* and *bud32* have a 40 bp overlap in the Native fusion plasmid as in the genome. In the Tandem separated plasmid, *kae1* and *bud32* expression was controlled by two independent *araS* promoters.

**Figure 8.**
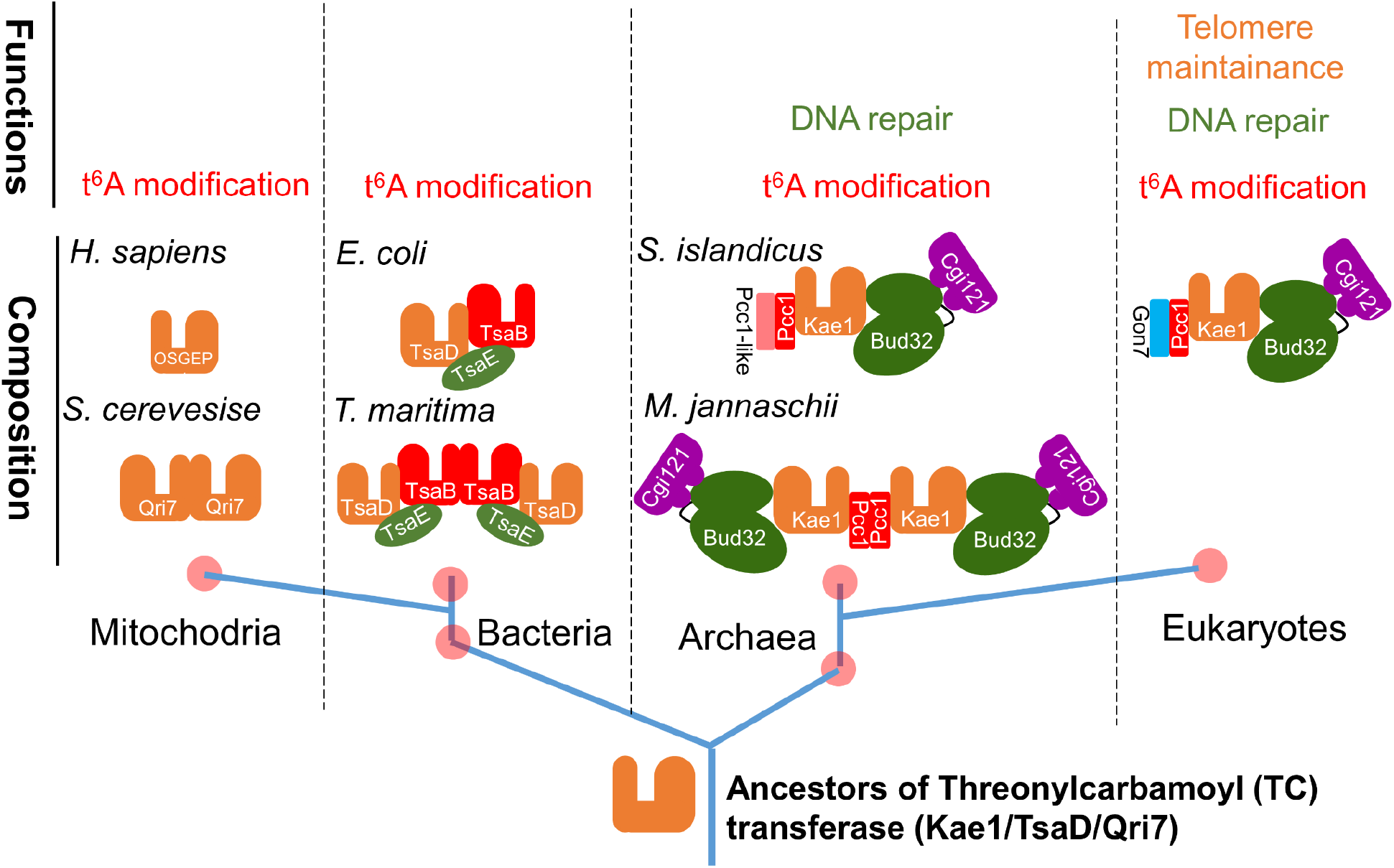
A proposed model for the evolution of TC-transferases and its complexes. The ancestral TC-transferase could be Kae1/TsaD, from which mitochondrial TC-transferase and bacterial TsaD originated functioning for TC transfer. Later, the ATPase Bud32/TsaE and the dimerization subunit Pcc1/TsaB joined to form the complexes comprising minimum components and essential for t^6^A modification. Meanwhile, Cgi121 joined into the initial complex of archaea-eukaryotes linkage to recruit tRNA. In the later stage, Pcc1-like might appear via a gene duplicate to stabilize the complex. Perhaps since Pcc1-like worked as a regulator in t^6^A modification, it exhibited the highest mutation rate among the subunits and evolved to Gon7 in eukaryotes. As Cgi121 and Pcc1-like (Gon7) were integrated into the complex, KEOPS gained additional functions other than t^6^A modification, such as DNA repair (HR repair) before the separation of archaea and eukaryotes. In eukaryotes, KEOPS also participates telomere maintenance which requires DNA recombination processes fulfilled by multiple HR repair proteins.

## Discussion

About sixty proteins are conserved in all cellular life, these include Sua5/TsaC and Kae1/TsaD/Qri7, the key enzymes in t^6^A modification (10,52). Kae1 in cooperation with Pcc1, Bud32, Cgi121, and eukaryotes specific fifth subunits Gon7, forms the KEOPS complex to catalyse t^6^A modification (46). Besides t^6^A modification, KEOPS complex also plays roles in HR repair and telomere maintenance in eukaryotes (13,21). Although it was shown that archaeal KEOPS functions in t^6^A modification, whether it has the fifth subunit and whether it has function beyond t^6^A modification is unclear. Here, we show that the paralog of Pcc1, Pcc1-like, is the fifth subunits of KEOPS in archaea and aKEOPS complex has essential role other cellular process besides t^6^A modification, presumably DNA homologous recombinational repair.

Although archaeal Pcc1, Kae1 and Bud32 was the minimal components for t^6^A modification activity *in vitro*, the eukaryotic fifth subunit Gon7 deletion strain also caused severe t^6^A modification defect in *S. cerevisiae*, indicating the critical role of the fifth subunit (12,46). Therefore, we searched for the distribution pf KEOPS subunits in archaea and identified a Pcc1 paralog, Pcc1-like, that is likely the functional homolog of Gon7. *In vivo* genetic analysis showed that *pcc1* and *pcc1-like* can neither be deleted, suggesting a non-redundant role of *pcc1-like. In vivo* and *in vitro* interaction analysis shows that Pcc1-like interacts with both Pcc1 and Kae1. Pcc1 inhibits the dimerization of Kae1-Pcc1 and stabilize the subcomplex *in vitro*, just like Gon7 (4,9). Additionally, bio-informational analysis shows that the Pcc1-Pcc1-like heterodimer interface resembles that of the eukaryotic Pcc1-Gon7 heterodimer. Interestingly, through sequence alignment and phylogenetic analysis, we found that the evolution rate of Pcc1 and Pcc1-like was higher than other KEOPS subunits and Pcc1-like lineage seems to have a closer relationship with Gon7 lineage than archaeal and eukaryotic Pcc1 in the phylogenetic tree (Fig. 1B), which supports that Pcc1-like is a functional homolog of Gon7. The fifth subunit Gon7 also functions in t^6^A modification and DNA repair in *S. cerevisiae* and *H. sapiens* (5,9,13,46). However, because its structure is partially disordered, the mechanism of Gon7 in these functions is still unknown (4,9). Structurally, Gon7, interacts with Pcc1 and stabilizes the KEOPS forming a 1:1:1:1:1 complex (4,9). The precise role of Pcc1-like/Gon7 in HR repair and t^6^A modification needs further study. To this end, Pcc1-like and aKEOPS as the whole complex from thermophilic archaea could be used as excellent structural and functional models for KEOPS study taking advantage of their stability.

We demonstrate that KEOPS has a t^6^A modification level-independent function that is essential in *S. islandicus*, a thermophilic crenarchaeon. To circumvent the possible essentiality of aKEOPS, we introduced a thermophilic bacteria-derived t^6^A modification system (45) to *S. islandicus* and performed genetic analysis of the KEOPS. We show that the bacterial t^6^A modification system is active in *S. islandicus* and can complement the t^6^A modification function of aKEOPS. Using this constructed knock-in strain TsaKI, we demonstrate that all the KEOPS subunits cannot be deleted and knock-down of Cgi121 leads to growth retardance. These results indicates that archaeal Cgi121 plays an important role in the cell besides t^6^A modification..

HR repair pathway is one of most complicated and the mechanism has not been fully elucidated. Among the DNA repair pathways of thermophilic archaea such as Sulfolobales and Thermococales, only the HR pathway is essential and most genes involved in HR cannot be knocked out (25,26,44,53). Considering this, the pathway in which KEOPS involved in *S. islandicus* is most likely HR repair. In halophilic euryarchaeon *H. volcanii, kae1, bud32*, and *cgi121* are essential for cell viability, whereas deletion of *pcc1* resulted in cells with higher nucleic acid content (28). The authors did not explain the cause of this observation, but we now assume that this could be due to the presence of higher genome copy number which may indicate that KEOPS is involved in DNA repair. *H. volcanii* is highly polyploid with approximately 20 copies of genomes per cell and is capable of origin-free replication depending on DNA recombination (54,55). Therefore, in addition to t^6^A modification, *H. volcanii* KEOPS might also contribute to this polyploidy species-specific, HR-dependent replication mode. In humans, knock-down of genes encoding KEOPS subunits induces the DNA damage response signalling and subsequent apoptosis (8). Therefore, involvement in DSB resection could be a conserved and ancestral function of KEOPS.

We found that overexpression of different subunits, their mutant, or co-expression of combinations of subunits had distinct effect on cell growth. The phenotypes resemble those in the yeast to some extent. It was reported that the activity residues of the kinase Bud32 plays a critical role in HR repair in *S. cerevisiae*, and the kinase activity of *S. cerevisiae* Bud32 was inhibited by Kae1 and activated by Cgi121 *in vitro* (3,13,56). These results suggest that there is a sophisticated regulatory mechanism of the KEOPS complex. The mechanism how aKEOPS participates in cellular processes such as HR and how this function is coordinated with the t^6^A modification will be an intriguing subject for further investigation.

A mystery of KEOPS is the molecular basis to explain why KEOPS has such seemingly un-related three functions, t^6^A modification, HR repair, and telomere maintenance in yeast. Up to now, mechanisms have been elucidated for the t^6^A modification of KEOPS except for the precise roles of Gon7 and Pcc1 in the process. Cgi121 was reported to be a tRNA 3’ CCA tail and ssRNA binding protein for the recruitment of tRNA to KEOPS during t^6^A modification (11). Interestingly, telomeres in most eukaryotes are composed of TT[A/T]GGG repeats, whose antisense strands and corresponding telomerase RNAs are CCC[T/A]AA repeats (57). Therefore, the nucleic acids CCA binding ability of Cgi121 may be a link among the three functions of eukaryotic KEOPS. Our study revealed that Cgi121 is the only subunit that binds to a long dsDNA individually, but not short ss/dsDNA (which might be not able to form a higher structure) containing a CCA motif. Strikingly, mutation at the tRNA binding residues of *Tko*Cgi121 led to a defect in long dsDNA binding ability, suggesting that the mechanisms of KEOPS in t^6^A modification and DNA repair are related (11). The evolutionary transition from circular to linear chromosomes, which harbour telomeric structures, brought more challenges to genome stability in eukaryotic cells. This could be the reason why multiple DNA repair proteins, especially classical HR proteins and KEOPS also participate in telomere maintenance.

Finally, based on studies on the composition and functions of the t^6^A modification systems, we proposed a model for the evolution of the KEOPS complex and other TC-transferase or its complexes (Fig. 7). The ancestral TC-transferase could be a single protein Kae1/TsaD, from which mitochondrial TC-transferase Qri7 and bacterial TsaD originated independently (46,58). Then, ATPase Bud32/TsaE and the dimerization module Pcc1/TsaB join to form an initial KEOPS complex, which contains the minimum components and is essential for t^6^A modification (45,46). Meanwhile, Cgi121 come into the initial complex of archaea-eukaryotes lineage to recruit tRNA (11). At a later stage, Pcc1-like appears via gene duplication, stabilizes the complex, and improves the efficiency of t^6^A modification (Fig. S6) (9). Finally, due to the accessory role of Pcc1-like in t^6^A modification, Pcc1-like exhibits the highest mutation rates among the subunits and evolved into Gon7 in eukaryotes. During the later stages as the additional subunits join, KEOPS gains functions other than t^6^A modification, such as HR repair, before the separation of archaea and eukaryotes. In eukaryotes, KEOPS is also involved in telomere maintenance which also needs a process of DNA recombination, like other HR repair proteins such as Mre11 and Rad50 (59,60).

## Acknowledgements

This work was supported by the National Key Research and Development Program of China (No. 2020YFA0906800), the National Natural Science Foundation of China (No. 31970546 and 31670061 to YS, 31900055 to QH, 31970119 to JN, and 31771380 to QS) and the State Key Laboratory of Microbial Technology. We would like to thank all the lab members of the CRISPR and Archaea Biology Research Centre for helpful discussions.

