## Supplemental Figures and Tables for "The archaeal KEOPS complex possesses a functional Gon7 homolog and has an essential function independent of cellular t^6^A modification level"

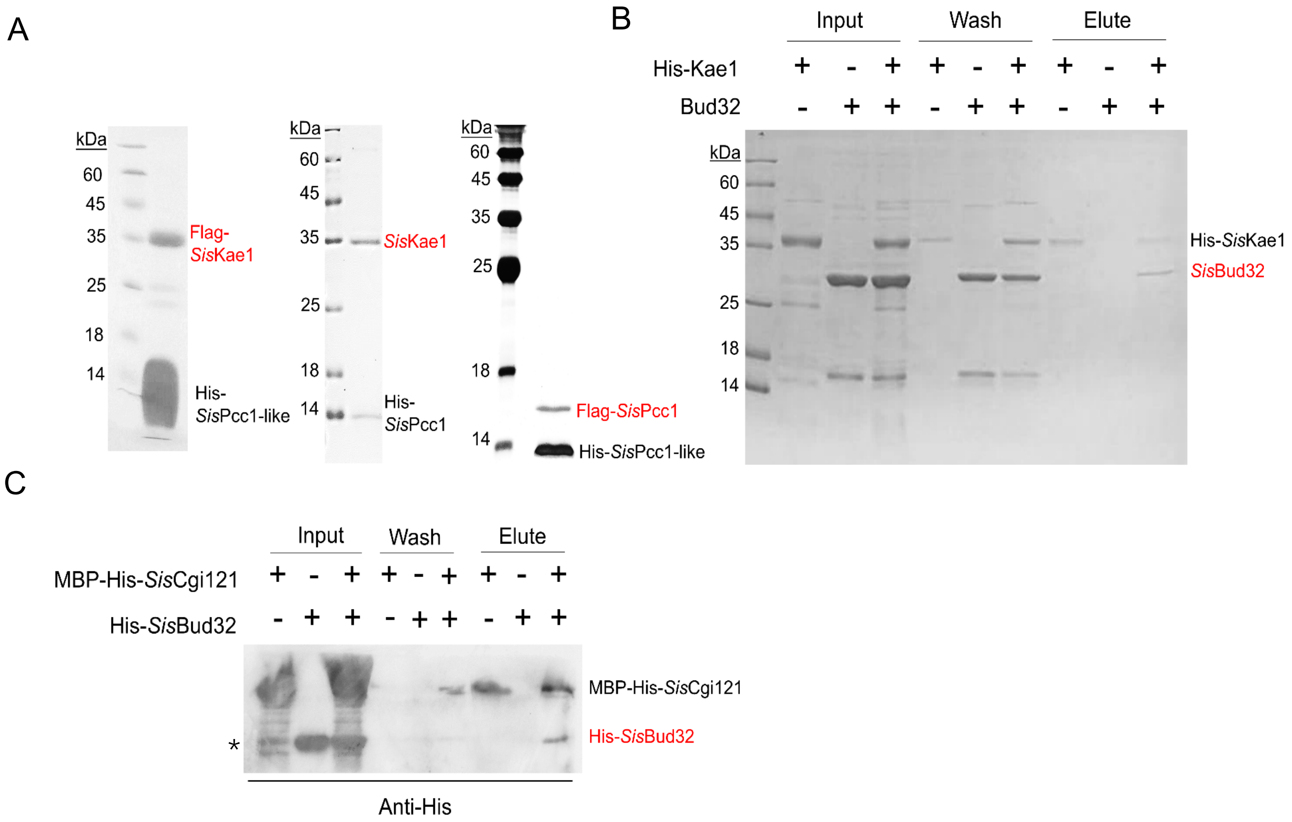


**Figure S1. Pair-wise interaction analysis of aKEOPS subunits by pull down.** (**A**) Assay using proteins expressed and purified from *E. coli*. The interactions between *Sis*Kae1 and *Sis*Pcc1-like, *Sis*Kae1 and *Sis*Pcc1, and *Sis*Pcc1 and *Sis*Pcc1-like were verified by the ability to pull down the target proteins (red) with the His-tagged proteins co-expressed. The final eluted samples from Ni-NTA beads were analyzed by SDS-PAGE. (**B**) *In vitro* pull down assay using purified His-*Sis*Kae1 as a bait and non-tagged *Sis*Bud32 as a prey. The samples were analyzed by SDS-PAGE and Coomassie blue staining. (**C**) *In vitro* pull down assay using MBP-His-*Sis*Cgi121 as a bait and His-*Sis*Bud32 with amylose resin. The samples were analyzed by Western blotting. In (B) and (C), all the proteins were expressed in *E. coli* individually and purified by affinity chromatography. The asterisk indicates a non-specific band in the MBP-His-SisCgi121 sample that cannot bind to amylose resin.

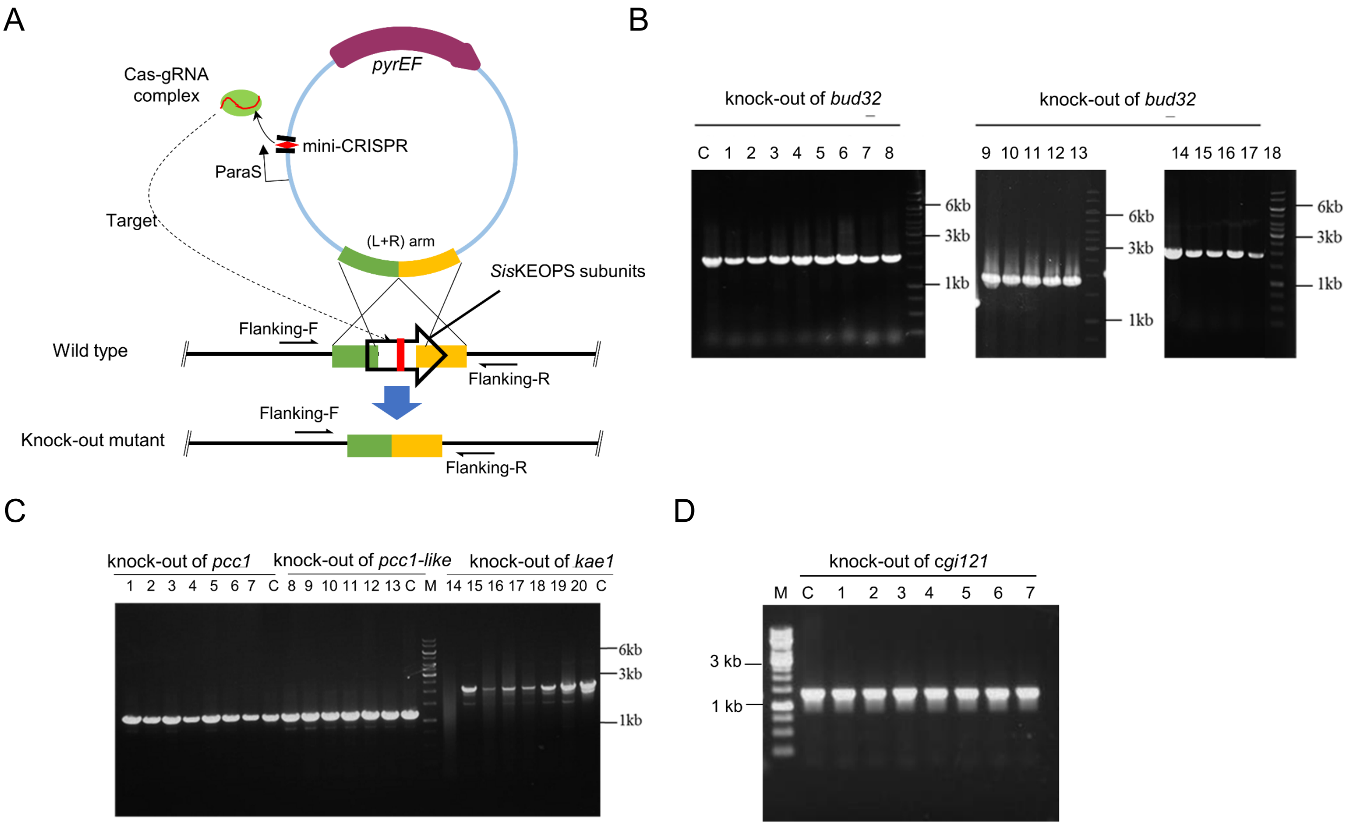


**Figure S2. PCR analysis of the colonies obtained in the *Sis*KEOPS gene knock-out experiments. (A)** Schematic of the gene knock-out strategy. **(B)** Screening for *bud32* knock-out colonies. **(C)** Screening for *pcc1*, *pcc1-like*, and *kae1* knock-out colonies **(D)** Screening for *cgi121* knock-out colonies. M, molecular size marker. C, control (E233S). The numbers indicate colonies picked from the plates after transformation.

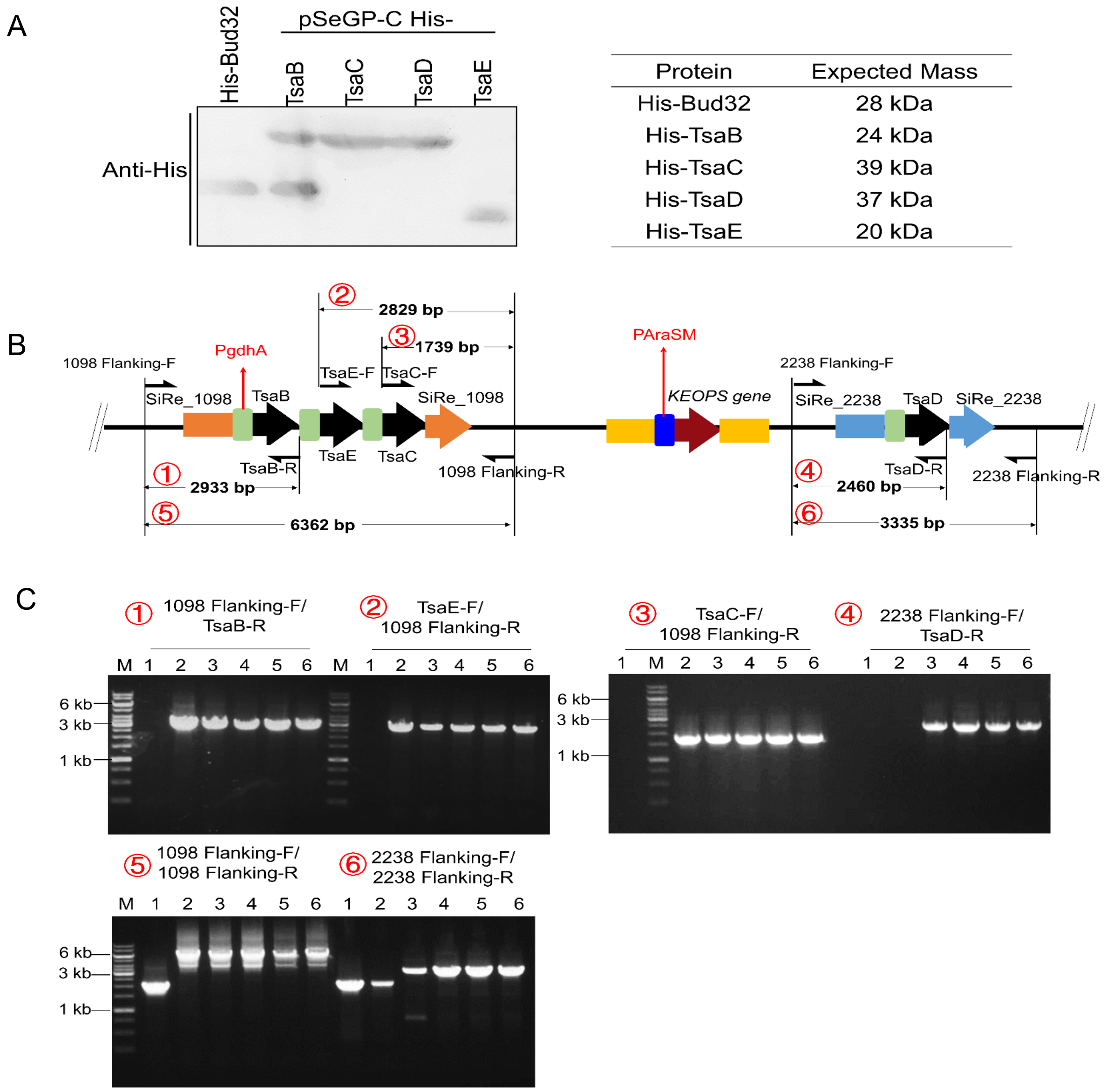


**Figure S3. Confirmation of protein expression of TmTsaB/C/D/E in the overexpression strains of E233S and verification of the insertion of *TmTsaB/C/D/E* in TsaKI. (A)** Western blotting analysis of strains over expressing each of TmTsaB/TsaC/TsaD/TsaE with C terminal His-tag (left) using the shuttle vector pSeGP. About $2.5\times10^{8}$ cells were collected. Purified His-tagged Bud32 was used as a positive control. Expected molecular mass of each protein is shown on right. (**B**) Schematic for the primers at the knock-in locus and the expected PCR products. The numbers in red indicated PCR products of their corresponding numbers in (B). (**C**) Analysis of the PCR products using primers at the *α-amylose* (*SiRe_1098*) and the *β-mannanase* (*SiRe_2238*) loci. Lanes 1, E233S; 2, E233S/*TmTsaBEC::amyα*; 3, TsaKI; 4, TsaKI/*$P_{araS}$24::kae1-bud32*; 5, TsaKI/*$P_{araS}$38::kae1-bud32*. 6, TsaKI/*$P_{araS}$38::cgi121*.

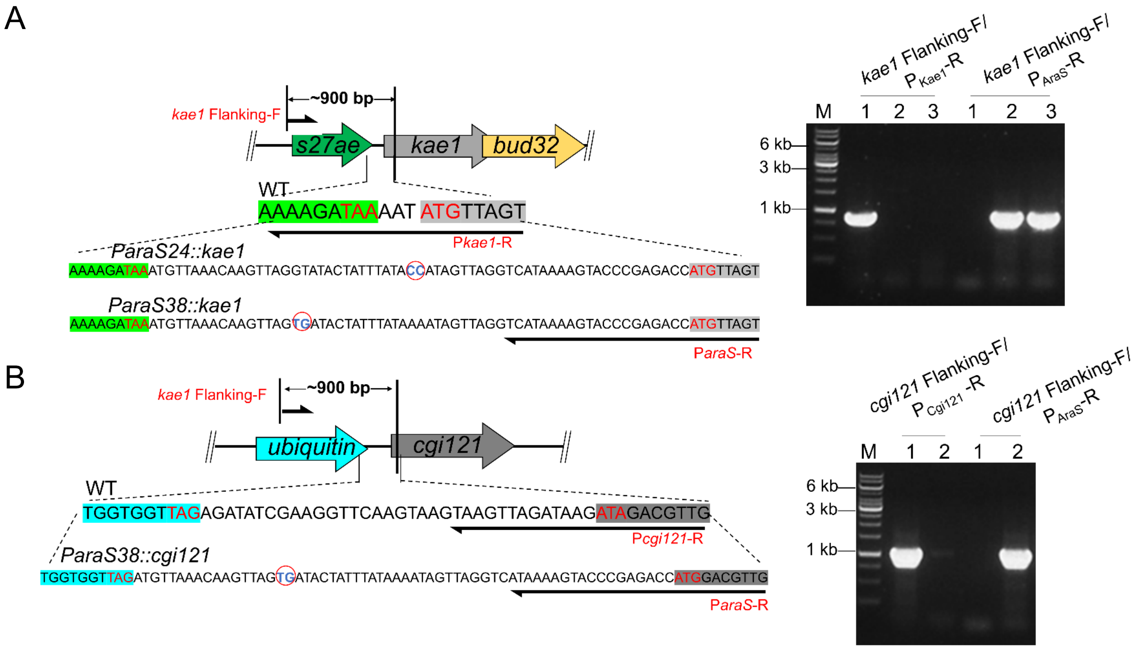


**Figure S4. PCR verification of the promoter replacement of the KEOPS genes in TsaKI.** (**A**) Analysis of PCR products using primers at the loci of *kae1* promoters (right). The schematic for the promoter replacement of *kae1-bud32* is shown on the left. The start and stop codons of *kae1* (or *cgi121* in D) and its upstream gene are indicated in red. The two nucleotides in red circular are mutations of the wild type arabinose promoters. 1, TsaKI; 2, TsaKI/$P_{araS}$*24::kae1-bud32*.; 3, TsaKI/$P_{araS}$*38::kae1-bud32*. (**B**) Analysis of the PCR products using primers at the loci of *kae1-bud32* promoters (right). The schematic for the promoter replacement is shown on the left. 1, TsaKI; 2, TsaKI/$P_{araS}$*38::cgi121*. M, marker.

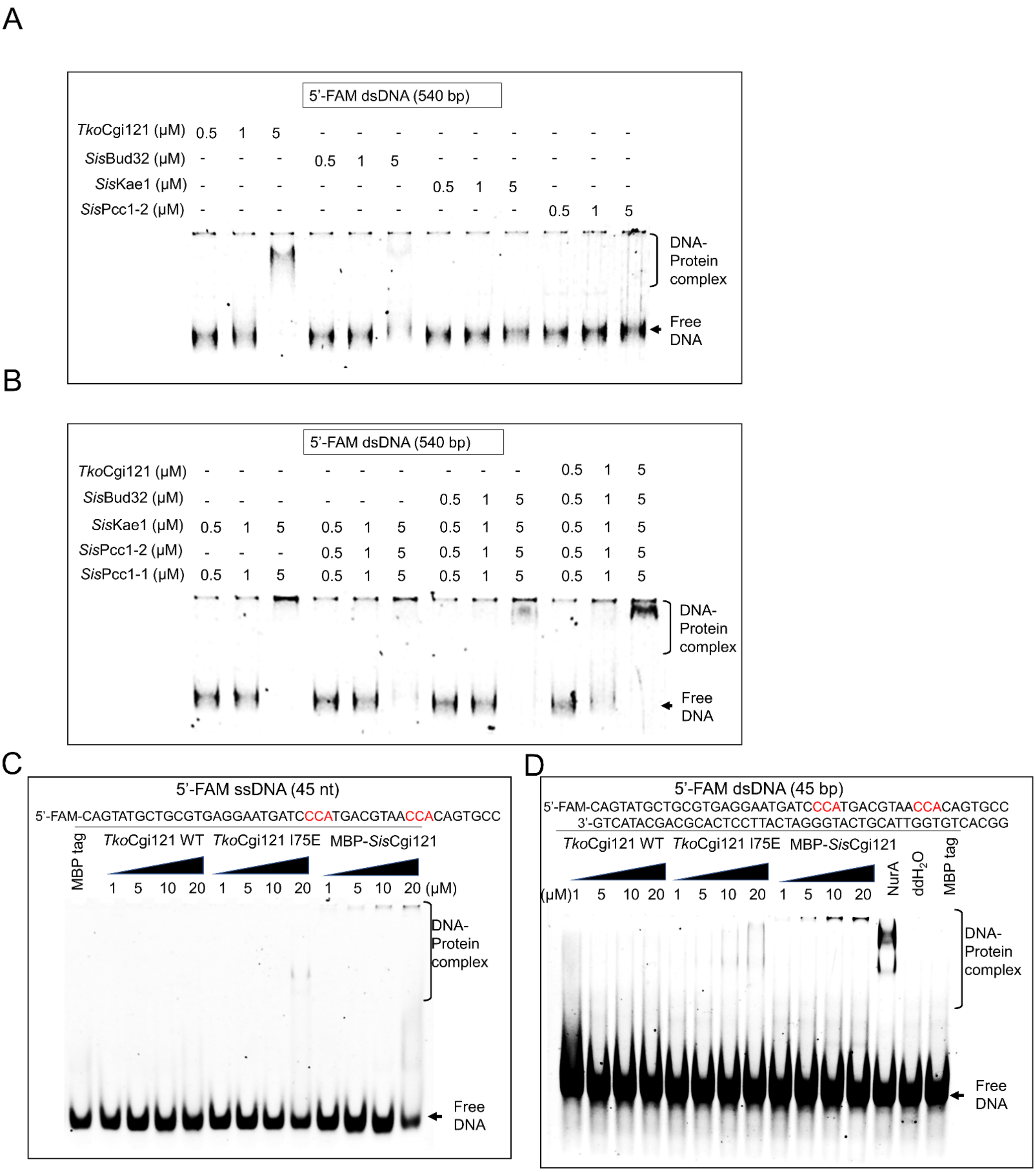


**Figure S5. Analysis of the DNA binding ability of archaeal KEOPS**. (**A**) Binding of individual subunit for dsDNA. (**B**) Binding of combinations of the subunits for dsDNA. FAM-labeled dsDNA (10 nM) was incubated with samples of the indicated subunits (0.5, 1, 5 μM) at 37C° for 30 min. (**C**) *Tko*Cgi121 (WT), the CCA-3' RNA binding site mutant *Tko*Cgi121 (I75E), and *Sis*Cgi121 for ssDNA binding. Purified Cgi121 (1, 5, 10, 20 μM) was incubated in a reaction mixture containing 10 nM of FAM-labeled ssDNA（45 nt, with two CCA motifs colored in red）at 37C° for 45 min. *Sis*SSB (5 μM) was used as a positive control and MBP-tagged *Sis*Pcc1 (20 μM) was used as negative control. (**D**) *Tko*Cgi121 (WT), *Tko*Cgi121 (I75E), and *Sis*Cgi121 for short dsDNA binding. Purified Cgi121 (1, 5, 10, 20 μM) was incubated in a reaction mixture containing 10 nM of FAM-labeled dsDNA (45 bp, with two CCA motifs colored in red) at 37C° for 45 min. *Sis*NurA (5 μM) was used as a positive control and MBP-tagged *Sis*Pcc1 (20 μM) was used as negative control. The samples were loaded in native PAGE gels. The gels were scanned using Amersham ImageQuant 800 after electrophoresis. The detailed reaction conditions are described in the Materials and Methods.

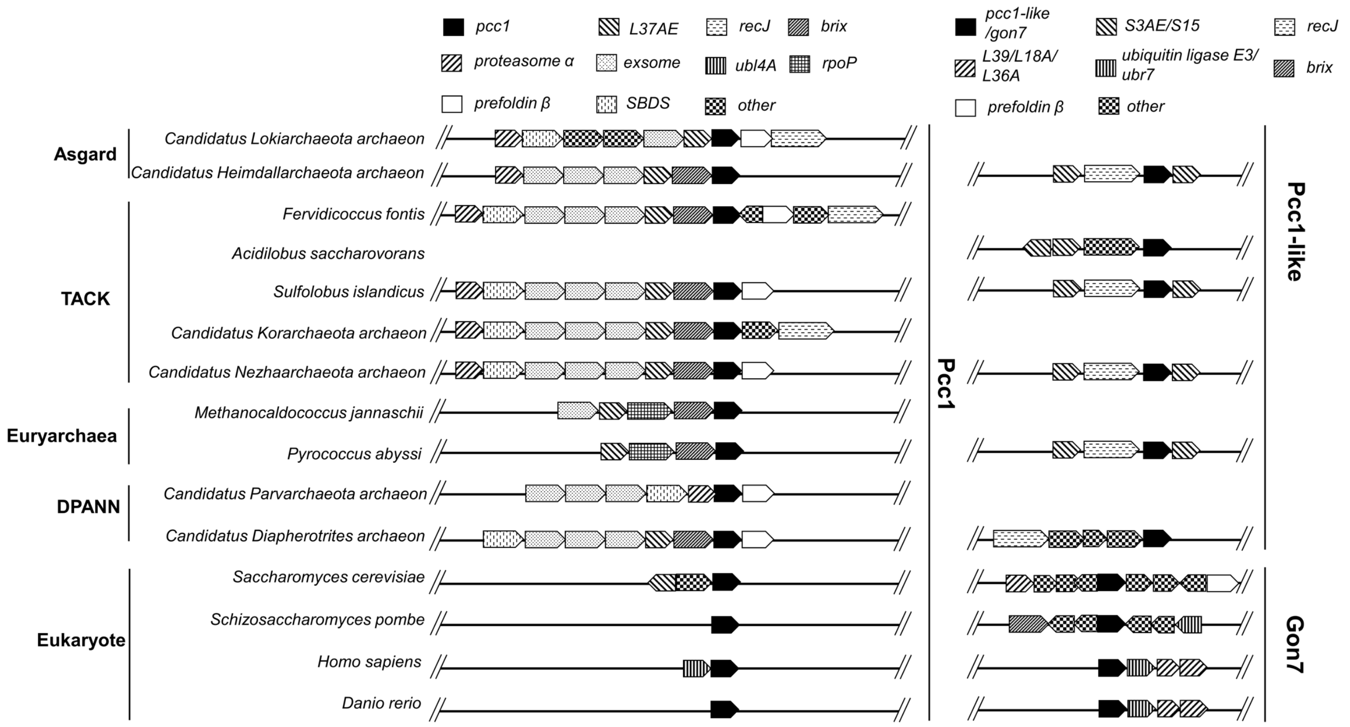


**Figure S6. Genome context analysis of *pcc1*, *pcc1-like*, and *gon7* in archaea and eukaryotes.** Representative species possessing one *pcc1* and two *pcc1* paralogs are shown for each superphylum of archaea together with several eukaryotic species. The *pcc1-like* occurred possibly via gene duplication of *pcc1* and was separated together with *recJ* during evolution.

**Table S1.** Strains used in this study

| Strain | Properties | Source or reference |
|---|---|---|
| *S. islandicus* E233S | *S. islandicus* REY15A Δ*pyrEF* Δ*lacS* | Peng *et al.*, 2009, 2012 |
| E233S/$P_{araS}$*24::kae1-bud32* | *kae1-bud32* promoter replaced with $P_{araS}$*24* in E233S | this study |
| E233S/$P_{araS}$*38::kae1-bud32* | *kae1-bud32* promoter replaced with $P_{araS}$*38* in E233S | this study |
| E233S/$P_{araS}$*24::cgi121* | *cgi121* promoter replaced with $P_{araS}$*24* in E233S | this study |
| E233S/*amyα::TmTsaB/TsaE/TsaC* | *TmTsaBEC* knocked in *amyα* (*sire_1098)* in E233S | this study |
| TsaKI | *TmTsaD* knocked in *mannanase* (*sire_2238*) in E233S/*amyα::TmTsaBEC* | this study |
| TsaKI/$P_{araS}$*38::kae1-bud32* | *kae1-bud32* promoter replaced with $P_{araS}$*38* in TsaKI | this study |
| E233S/$P_{araS}$*38::cgi121* | *cgi121* promoter replaced with $P_{araS}$*38* in TsaKI | this study |

**Table S2.** Vectors used in this study.

| Vectors | Properties or usages | Source or reference |
|---|---|---|
| pGE | *Sulfolobus*-*E. coli* shuttle vector containing mini-CRISPR and *pyrEF* for CRISPR-Cas based gene editing | Li *et al.*, 2016 |
| pGE-*kae1*mg-KD | knock down of *kae1* with multi-gRNA | this study |
| pGE-*bud32*mg-KD | knock down of *bud32* with multi-gRNA | this study |
| pGE-*pcc1*sg-KD | knock down of *pcc1* with single-gRNA | this study |
| pGE-*pcc1-like*sg-KD | knock down of *pcc1-like* with single-gRNA | this study |
| pGE-*kae1*-KO | knock out of *kae1* with single-gRNA | this study |
| pGE-*bud32*-KO | knock out of *bud32* with single-gRNA | this study |
| pGE-*kae1-bud32*-KO | knock out of kae1-*bud32* with multi-gRNA | this study |
| pGE-*cgi121*-KO | knock out of *cgi121* with single-gRNA | this study |
| pGE-*pcc1*-KO | knock out of *pcc1* with single-gRNA | this study |
| pGE-*pcc1-like*-KO | knock out of *pcc1-like* with single-gRNA | this study |
| pGE-*cgi121*M52E | construction of *cgi121* mutant M52E with single-gRNA | this study |
| pGE-*cgi121*I64E | construction of mutant *cgi121* mutant I64E with single-gRNA | this study |
| pGE-*TmTsaBDEC*-KI | *tsaBDEC* knock in at *amyα* locus with single-gRNA | this study |
| pGE-*TmTsaD*-KI | *TmtsaD* knock in at *mannanase* locus with single-gRNA | this study |
| pSeSD | *Sulfolobus*-*E. coli* shuttle vector containing *araS*-SD promoter and MCS for proteins expression in E233S | Peng *et al.*, 2012 |
| pSeSD-His-*Sis*Kae1 | expression of C-terminal 6×His tagged *Sis*Kae1 | this study |
| pSeSD-His-*Sis*Bud32 | expression of C-terminal 6×His tagged *Sis*Bud32 | this study |
| pSeSD-His-*Sis*Bud32D134A | expression of C-terminal 6×His tagged *Sis*Bud32D134A | this study |
| pSeSD-His-*Sis*Cgi121 | expression of N-terminal 6×His tagged *Sis*Cgi121 | this study |
| pSeSD-His-*Sis*Pcc1 | expression of C-terminal 6×His tagged *Sis*Pcc1 | this study |
| pSeSD-His-*Sis*Pcc1-like | expression of C-terminal 6×His tagged *Sis*Pcc1-like expression | this study |
| pSeSD-*Siskae1*-(His)*bud32* | expression of *Siskae1-bud32* operon with bud32 being tagged with C-terminal 6×His | this study |
| pSeSD-2ParaS | containing two *araS* promoter for protein expression | this study |
| pSeSD-2ParaS-*Sis*Kae1/His-*Sis*Bud32 | Co-expression of *Sis*Kae1 and C-terminal 6×His tagged *Sis*Bud32 | this study |
| pSeSD-2ParaS-His-*Sis*Kae1/ His-*Sis*Bud32 | Co-expression of C-terminal 6×His tagged *Sis*Kae1 and C-terminal 6×His tagged *Sis*Bud32 | this study |
| pSeSD-2ParaS-His-*Sis*Cgi121/His-*Sis*Bud32 | N-terminal 6×His tagged *Sis*Cgi121 and C-terminal 6×His tagged *Sis*Bud32 coexpression vector | this study |

| | | |
|---|---|---|
| pSeGP | *araS-SD* replaced with P*gdhA* in pSeSD | this study |
| pSeGP-His-*Tm*TsaB | constitutive expression of *Tm*TsaB | this study |
| pSeGP-His-*Tm*TsaC | constitutive expression of *Tm*TsaC | this study |
| pSeGP-His-*Tm*TsaD | constitutive expression of *Tm*TsaD | this study |
| pSeGP-His-*Tm*TsaE | constitutive expression of *Tm*TsaE | this study |
| pET22b-His-*Sis*Cgi121 | expression of the N-terminal 6×His tagged *Sis*Cgi121 | this study |
| pET22b-His-*Sis*Cgi121 | expression of C-terminal 6×His tagged *Sis*Cgi121 | this study |
| pET15b-*Sis*Kae1 | expression of no tagged *Sis*Kae1 | this study |
| pET15b-*Sis*Bud32 | expression of no tagged *Sis*Bud32 | this study |
| pET15b-His-*Sis*Bud32 | expression of N-terminal 6×His tagged *Sis*Bud32 | this study |
| pRSFDuet1-His-*Sis*Kae1 | expression of N-terminal 6×His tagged *Sis*Kae1 | this study |
| pRSFDuet1-Flag-*Sis*Bud32 | N-terminal Flag tagged *Sis*Bud32 expression vector | this study |
| pRSFDuet1-His-*Sis*Pcc1 | N-terminal 6×His tagged *Sis*Pcc1 expression vector | this study |
| pRSFDuet1-His-*Sis*Pcc1-like | N-terminal 6×His tagged *Sis*Pcc1-like expression vector | this study |
| pRSFDuet1-His-*Sso*Cgi121 | expression of N-terminal 6×His tagged *S. solfataricus* Cgi121 | this study |
| pRSFDuet1-His-*Tko*Cgi121 | expression of N-terminal 6×His tagged *T. kodakarensis* Cgi121 | this study |
| pRSFDuet1-His-*Tko*Cgi121I75E | expression of N-terminal 6×His tagged *Tko*Cgi121(I75E) | this study |
| pRSFDuet1-His-*Sis*Pcc1/*Sis*Kae1 | co-expression of N-terminal 6×His tagged *Sis*Pcc1 and no tagged *Sis*Kae1 | this study |
| pRS Duet1-His-*Sis*Pcc1-like/Flag-*Sis*Kae1 | co-expression of the N-terminal 6×His tagged *Sis*Pcc1-like and N-terminal Flag tagged *Sis*Kae1 | this study |
| pRSFDuet1-Flag-*Sis*Pcc1/His-*Sis*Pcc1-like | co-expression of the N-terminal Flag tagged *Sis*Pcc1 and N-terminal 6×His tagged *Sis*Pcc1-like | this study |
| pMALc2X-MBP-His-*Sis*Cgi121 | co-expression of the N-terminal MBP and C-terminal 6×His tagged *Sis*Cgi121 | this study |
| pMALc2X-MBP-His-*Sis*Pcc1 | co-expression of N-terminal MBP and C-terminal 6×His tagged *Sis*Pcc1 | this study |

**Table S3.** Oligonucleotide and promoter DNA sequences used in this study.

| Oligonucleotides | Sequences (5'to3') |
|---|---|
| *kae1*-KO-Spacer | GGCGGAAATACAATCATAACTACCTTCTATAAAGGGAGGT |
| *bud32*-KO-Spacer | GCGGTGATCTAACAACTAACAATCTCATCCTAAGTTCTATA |
| *kae1-bud32*-KO-Spacer1 | ACTATGAAAGGACAATATTAGAGGCTAAGATAATTTATAC |
| *kae1-bud32*-KO-Spacer2 | GGCGGAAATACAATCATAACTACCTTCTATAAAGGGAGGT |
| *kae1-bud32*-KO-Spacer3 | AGGAATATCTACCTCGTCTACTCTCCATCTAGGTCTTAT |
| *cgi121*-KO-Spacer | ACTATAATACAACTAGAAATAAGATAAAGAGTTCAACTAT |
| *pcc1*-KO-Spacer | ATAAAAACGAATTACAGGATATAATATATGATTCGATAAT |
| *pcc1-like*-KO-Spacer | ATTCAAACCAAGGTTGAAGGCAAAGAGTTAGAGATTGTAA |
| *kae1*-KD-Spacer1 | TCAGAATGTATACTTTCTAAAGATCTTAGAAATATATGAA |
| *kae1*-KD-Spacer2 | ATATCGTTATTTATAGAACTTAGGATGAGATTGTTAGTTG |
| *kae1*-KD-Spacer3 | CACTGCAGGTACATTCACATCGTTTTTAAGCGCAGTATAA |
| *bud32*-KD-Spacer1 | GGTCTTTGGCCTCAGTTGTTAAATACCCTATTTCAATATG |
| *bud32*-KD-Spacer2 | CTACTGCTATATAATTAATATCATTTATACTAATATTAGC |
| *bud32*-KD-Spacer3 | TATCCCTTTCGTTTGCCAATATGTAGGGTGGTTGATCTTT |
| *pcc1*-KD-Spacer | GGGATTTCTTAATTTTTACATATTTAGTATCTATCTTTTC |
| *pcc1-like*-KD-Spacer | TCGTTTGGCTTTGTAATTCTCATCTTTATTACAATCTCTA |
| *kae1-bud32*-PE-Spacer | AAGAAAAGATAAAATATGTTAGTACTGGGTATCGAATCTA |
| *cgi121*-PE-Spacer | TATCTAACTTACTTACTTGAACCTTCGATATCTCTAACCA |
| *cgi121*M52E/I64E-Spacer | TACTTCTCCTATTACCATATGAACAAATAAAGGATGCATT |
| 1098-KI-Spacer | TTATTCTCAAGCTCTATTGGTAGCCATTTGAGAAATTCTA |
| *2238*-KI-Spacer | ATGCACCATCAGCACCGTAAACTAACGCTCCCCTCAGTAT |
| $P_{araS}$*24* | ATGTTAAACAAGTTAGGTATACTATTTATACCATAGTTAGGTCATAAAAGTACCCGAGACC |
| $P_{araS}$*38* | ATGTTAAACAAGTTAGTGATACTATTTATAAAATAGTTAGGTCATAAAAGTACCCGAGACC |
| $P_{gdhA}$(with 30bp *gdhA* leader) | (-556)AACACTAATGAGAAAGT……CCGGGAAAACGAATTTATATTG**ATG**GAAGAAGTTCTTAGTTCGAGTCTGCAT (+30) * |
| 5'-FAM labled ssDNA | CAGTATGCTGCGTGAGGAATGATCCCATGACGTAACCACAGTGCC |
| *kae1*-qPCR-F | CTCGGAGTAGGAATAGCAAAAGATCA |
| *kae1*-qPCR-R | CGTTTTAGTAGATCTCCAGGCTTCAT |
| *bud32*-qPCR-F | GAGGCTAAGATAATTTATACTGCGCTT |
| *bud32*-qPCR-R | CCTTAACTATCTCCCCTTCTATATATTC |

| | |
|---|---|
| *cgi121*-qPCR-F | GTCAAGTCTTACCTATCGTACCGTTT |
| *cgi121*-qPCR-R | CGGTAATAGGAGAAGTAAGAAAAACATAG |
| *pcc1*-qPCR-F | GTGAAGATAGAGATTTCAATTTATCCAGATAAT |
| *pcc1*-qPCR-R | CCTAGCTCTTGTAATTGATGGTGCA |
| *pcc1-like*-qPCR-F | TTACCAGAAGGAATGTCAATTCAAACCA |
| *pcc1-like*-qPCR-R | TTATAGCATCAAATGACGATTGAAGTGC |
| *1715*-qPCR-F | TAACGGTGTCAGAGATAACCTACAC |
| *1715*-qPCR-R | TGTTATGTCCTTTACTCTTATACCTACC |
| *tbp*-qPCR-F | CAACAGTTACGTTAGAGCAAAGTTTGG |
| *tbp*-qPCR-R | GTAACCTTGGGCTGTTCTAATCTGA |

* The numbers in the brackets indicate the position of the last nucleotide at the upstream (-) or downstream (+) of the start codon (in red).
